# Petabase-scale Papillomavirus Discovery

**DOI:** 10.64898/2026.04.21.719858

**Authors:** Jessica Y. Q. Shen, Anton Korobeynikov, Rayan Chikhi, Artem Babaian

## Abstract

Freely available nucleic acid sequencing databases have accumulated to a vast archive of genetic diversity, in excess of 50 petabase-pairs from tens of millions of experiments. Together, these data constitute a digital survey of Earth’s genome. However, the richness of biological information contained within these repositories remains largely unexplored, in large part owing to the technical challenges of analyzing petabytes of data. Recently, *Logan* completed the sequence-assembly and compression of 27 million sequencing libraries from the Sequence Read Archive (SRA), and here, we systematically search Logan-SRA to reveal the global diversity of the DNA-based Papillomaviruses (PVs).

In a single ∼10-hour alignment-based search against the *Logan* assemblage, we independently re-identified 65% of the 992 PVs recorded within the NCBI Virus database, a body of work representing over five decades of PV characterization. We further expand the diversity of PVs by 34%, identifying 383 novel PV types spanning 105 associated host species, including taxa with no previously associated PVs, such as rhinoceros, voles, and grey foxes. Through integration of virus phylogeny, sample geography, and ecological metadata, we show that novel PV discovery is not directly proportional to sampling effort, and there are significant hotspots of PV biodiversity in East Africa and South America, and that undersampled biomes can yield disproportionately more novel PVs.

Public sequencing repositories contain vast, unrealized biological information that is now accessible through advanced computational infrastructure. Here we lay the foundations for the efficient analysis of petabase-scale datasets for DNA virus discovery, with applications to uncovering all genetic diversity.

## INTRODUCTION

### Using digital archives to map genetic diversity

From fundamental clinical research, to expansive ecological surveys, the use and volume of high-throughput sequencing data has grown explosively over the last two decades. The combined scientific endeavours of the biological research community have led to the accumulation of a freely-available genetics heritage, spanning the vast diversity of life on our planet^1^.

The Sequence Read Archive (SRA) is the major sequence data-repository, representing over thirty-eight million publicly available sequencing libraries^2^. These libraries capture genetic information across time, geographic-location, source organisms, tissues, and environments, which together can be viewed as a single system, a digital ‘Earth genome’. The vast volume of predominantly short-read data is measured at the petabase scale (10^15^ base pairs), creating both significant hardware and algorithm challenges to its analysis, and opportunities for computational innovation.

Over the last decade, a new generation of computational tools have begun to solve the challenge of analyzing vast genetics databases with varying approaches. The NCBI Sequence Taxonomy Analysis Tool (STAT) is a pre-computed kmer index of known taxa genomes across the SRA, processing 27.9 Pbp over 5 years (as of 2021)^3^. Pebblescout, enables near instant nucleotide k-mer based search of over 3.7 Pbp of SRA and demonstrated the identification of drug-resistant *Candida auris*, in 129 SRA runs, relative to only 19 runs from a previous meta-data filtered study, highlighting the how metadata-agnostic or incidental nucleic acids are informative to broad ecological surveys^4^. Most recently MetaGraph expanded to k-mer alignment over an annotated *de Brujin* graph of over 5 Pbp of raw sequence data and used this to track antimicrobial resistance across 241,384 human gut microbiomes^5^.

In contrast, *Serratus*, a cloud-based translated-nucleotide search tool screened 5.7 million SRA datasets (10.2 Pbp) for the RNA-dependent RNA polymerase hallmark gene, but required 11 days of compute. This yielded >130,000 novel RNA viruses, and expanded the known diversity of RNA viruses by roughly an order of magnitude^6^. *Serratus* represents the first petabase-scale virus discovery effort, and highlights the trade-off for virus discovery between speed (kmer-based methods for exact or near-exact genome match) and sensitivity (for diverged protein sequences)^6^. Finally, these approaches are best when they can be combined, for example, the *Serratus* and *Pebblescout* platforms were instrumental in uncovering the diversity of an entirely novel class of viroid-like RNAs, Obelisks, over public sequencing datasets^7–9^. These genetic elements have no homologous relationships to any known lifeforms, and their molecular biology and ecology are still unknown^9^.

Collectively, these advancements establish public repositories not as mere “archival data”, but as a rich source of biodiversity, albeit with limited infrastructure to analyze it.

### Logan unlocks petabase-scale sequence analysis via large-scale assembly

Sequence assembly is another modality for compressing raw sequence data, converting fragmented reads into longer, more informative contigs, but it is computationally expensive.

To compress the SRA for efficient sequence search, we recently constructed *Logan*, a pan-SRA sequence assembly using cloud-computing of 96% of the SRA into contigs on a per-library basis. The freely-available *Logan* (v1.1) consists of 26,788,835 assembled accessions, or 48.2 × 10^15^ bases of data, capturing 88% of the SRA by bases sequenced^10^. Due to the compression and simplification of short reads, *Logan* offers the first opportunity to explore the untapped ‘digital ecology’ of the near totality of the SRA using sensitive alignment-based methods.

While there are many avenues of inquiry that could be explored via *Logan*, virus discovery stands to greatly benefit from this development. Traditional computational virus discovery studies tend to rely on targeted library sampling or sequence-generation, and are biased in terms of geography, host access, and study design. In contrast, viral sequences are commonly incidentally captured as ‘by-catch’ in libraries across a wide variety of studies, given their ubiquitous prevalence^11–13^. Thus, the SRA likely contains a vast, largely under-surveyed reservoir of viral diversity, which can now be explored systematically using *Logan*.

### A Papillomaviridae-based blueprint for pan-SRA DNA virus discovery

Here, we propose a framework for mining the *Logan* assemblage for novel Papillomaviruses (PVs). The *Papillomaviridae* (PV) family is an ideal candidate for DNA virus discovery within *Logan*. These non-enveloped, double-stranded DNA viruses are ancient, thought to have arisen around four hundred million years ago, and are presumed to infect the epithelial tissue of all major jawed vertebrate species^14,15^. All known PVs contain the core genes E1, E2, and L1, L2 which control early viral gene expression and replication, and form late structural components respectively^16^. PVs genomes can also contain accessory genes which can modulate the host cell through a wide variety of virus-host interactions to promote viral fitness, such as direct binding or sequestering of core cell-cycle control factors^17,18^.

Within the known range of hosts, PVs are commonly found within the virome of normal, immunocompetent individuals, and are often considered ‘commensals’ rather than being pathogenic^19,20^. In some cases, either mutation of the viral genome or integration into the host genome can cause PVs to become oncogenic, with the potential of this interaction depending on the presence of specific accessory genes, such as E6 or E7, in humans.^21^

PVs species biodiversity, called “types” within this viral family, are curated using the conserved hallmark gene, L1^22^. This gene encodes the major capsid protein of the PV, which contains the jellyroll fold, a structurally conserved motif consisting of two opposed four-stranded anti-parallel beta sheets arranged in a ‘sandwich’, with variable loops interspersed (Figure 1A)^23^. This structure is conserved across all PVs (Figure 1B) and is considered one of the most deeply shared structural features of viruses, encompassing more viral diversity than the RNA-dependent RNA polymerase (RdRp)^24,25^. A PV type is defined as a novel if its L1 is more than 10% diverged at the nucleotide level (< 90% nucleotide identity, nt id) from any known type^22,26^. Given the deep conservation of the L1 hallmark fold, and sequence-based taxonomic framework, makes PVs a well-defined viral family for a pan-SRA search. As such, the approach presented here for PVs represents a pilot study for the global survey of jellyroll fold viral diversity across the SRA.

**Figure 1.**
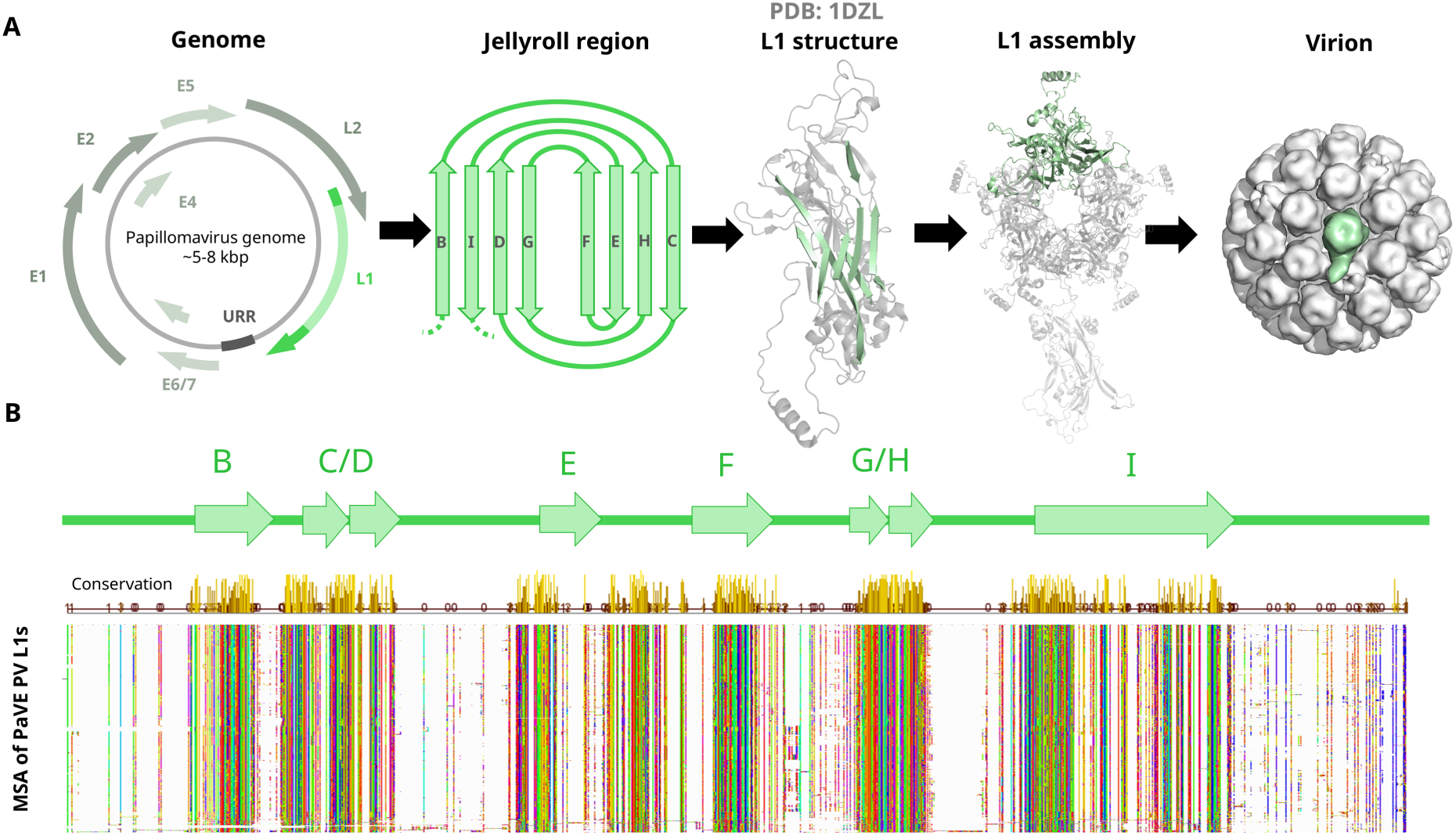
Papillomavirus genome organization and conservation of the L1 jellyroll fold. A. Prototypic arrangement of the papillomavirus (PV) genome. The PV genome consists of 6–8 open reading frames (green arrows) and an untranslated regulatory region (dark grey) arranged with conserved synteny. Core genes (dark sage), including the L1 major capsid protein (bright green) and its associated jellyroll region (light green), are found in all known PVs. The presence of accessory genes (light sage) varies between PV types. The PV capsid is composed of 72 self-assembling pentamers of the L1 protein, with the jellyroll region (represented here in both 2D topology and 3D structure) forming the principal structural fold. Protein and capsid structures retrieved from the HPV 16 L1 structure (PDB: 1DZL) and visualized in PyMOL^37,38^. B. The jellyroll fold is well-conserved across known PVs. A multiple sequence alignment (MUSCLE5 ‘super5’) from annotated PV L1 sequences in the Papillomavirus Episteme (PaVE, n = 614), clustered at 90% amino acid identity (n = 548, *Supplementary File* S1)^27,78^. Jellyroll beta-strands (B, C, D, E, F, G, H, I) were manually annotated based on the HPV 16 L1 structure (PDB: 1DZL) and residue positions in PyMOL^37,38^. Sequences are ordered by Jalview neighbour-joining tree (BLOSUM62) and visualized with the Taylor colour scheme^106^. Positional conservation is shown in the chartreuse histogram at the top of the alignment.

Human PVs are clinically relevant and therefore the focus of decades of research, however it is likely that significant portions of total PV diversity remain uncharacterized. Consider that the Papillomavirus Episteme (PaVE), a curated resource of PV genomes and meta-data contains 461 distinct human PV type genomes (<90% L1 nt id). However, of the 60,000+ other non-human vertebrate species, only 416 additional PV types are recorded, <1% of vertebrate species have a characterized PV^27^. This imbalance is likely due to under sampling of non-human host taxa, which is also reflected in the SRA. For instance, *Actinopterygii* (all ray-finned fish, taxid:7898) account for 724,194 SRA runs, while *Homo sapiens* (taxid:9606) are represented in 6,809,500 SRA runs [2026-04-08].

In a recent survey of over 127,000 SRA libraries, Buck et al. (2024), uncovered extensive novel diversity among small DNA tumor viruses, including novel PVs, polyomaviruses, parvoviruses, and previously unrecognizable viral gene classes in conjunction with these genomes, demonstrating the substantial DNA virus diversity still yet to be discovered within public sequencing data^28^.

The SRA, containing data from millions of sequencing studies without the express goal of PV discovery, yet many may harbor uncharacterized PVs which we aim to discover. To accomplish this, we analyze 26.8 million assemblies to identify 2.76 million PV contigs across 985,583 sequencing libraries, describing 383 novel PV types, spanning 105 distinct associated host species from all continents.

## RESULTS

### The SRA Contains Hundreds of Unidentified Papillomaviruses

#### Both known and novel PVs can be rapidly retrieved across the Sequence Read Archive

To establish a baseline of the known biodiversity of PVs, we extracted all *Papillomaviridae* nucleotide sequences in NCBI Virus (date accessed: 2024-06-11). Since PV genomes are circular and ORFs can be interrupted by linear breakpoints or genomic-insertions, we performed a stop-stop six-frame translation (*EMBOSS*) of the nucleotide sequences, and filtered and trimmed ORFs to match the known domains E1, E2, L1, or L2 with Pfam HMM models (Methods). This yielded 57,107 unique PV ORFs (12,463 E1; 12,208 E2; 20,870 L1; 11,566 L2), which clustered into 4,200 ORFs at 90% amino acid identity (877 E1, 915 E2, 1,452 L1, 855 L2), in a sequence set we call PVDB1.

To measure global papillomavirus biodiversity, we performed two iterative searches against the *Logan*-SRA assemblage^10^. First, we searched *PVDB1* (n = 4,200) using *DIAMOND blastp* against *Logan*-SRA v1.0 (26,785,902 sequencing datasets – 2024-07-09), extracted Logan PV matches and updated NCBI Virus sequences which we re-clustered to create *PVDB2*. Second, we searched *PVDB2* (n = 30,705) using *DIAMOND blastp* against *Logan*-SRA v1.1 (26,788,829 sequencing datasets – 2025-02-04). This returned 2,763,517 contigs across 985,583 (3.67% of SRA) datasets containing at least one significant PV-ORF (e-value < 1e^-10^).

Since L1 has been used as the taxonomic hallmark gene for PV in the literature, we next focused on identifying representative full-length L1 sequences, unless otherwise indicated^17,22,26,29,30^. 14,048 full-length PV L1 sequences were gathered from 7,433 libraries. Since *Logan* retains *de Brujin* graph assembly information in contig headers, we re-analyzed 18,642 *Logan* assemblies with fragmented, but significant L1 hits using a graph-aware HMM alignment tool, *Pathracer*, and recovered an additional 7,275 (51.7%) full-length L1 sequences^31^. This yielded a total, 21,319 full-length L1-containing contigs (Figure 2).

**Figure 2.**
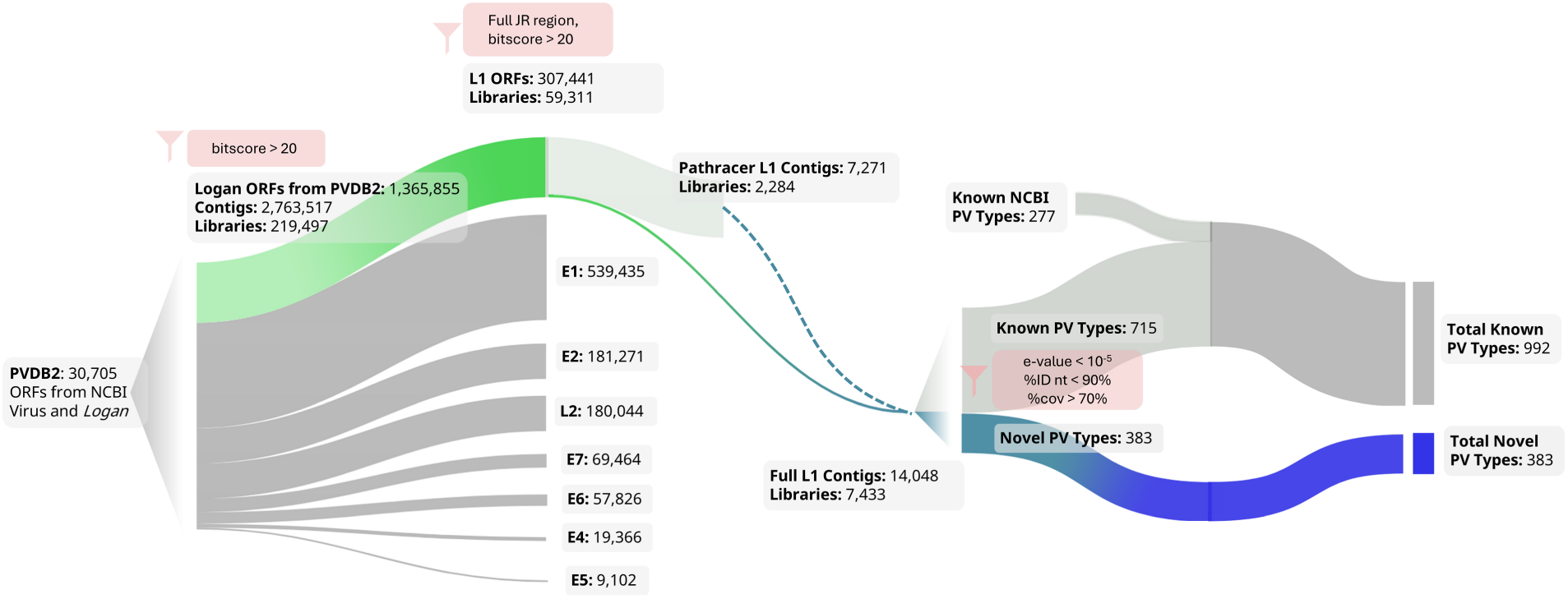
Discovery pipeline for novel papillomaviruses from *Logan*. Sankey diagram illustrating the filtering of PV-homologous sequences retrieved from *Logan*. Cutoff values and filtering criteria are indicated in pink at each step. Open reading frames (ORFs) with homology to PV genes were identified with HMMs from PVDB2 using a bitscore threshold of >20, with L1 sequences further filtered to retain only those spanning the full jellyroll region. Graph traversal using PathRacer was used to retrieve additional full L1 contigs from libraries with partial hits (dashed teal line)^31^. To reduce redundancy and classify PVs at a ‘type’ level, full L1s were clustered at 90% nucleotide identity to generate type-representative sequences

#### The SRA contains unexplored PV diversity

To measure L1 diversity in the SRA, the 21,319 full-length L1 nucleotide sequences were clustered at 90% identity into 1,097 representative PV L1 types. The metadata of each centroid was used to represent the putative host group of the cluster as above. To classify if a PV type has been described, we performed a *blastn* search of the centroid sequence against the *nt* database, a potentially more comprehensive but less curated database relative to NCBI Virus. To designate “Known PV Types”, 65% (714/1,097) of the PV types matched a *nt* sequence at ≥ 90% nucleotide similarity and 70% query coverage. In contrast, 35% types (383/1,097) were < 90% nt similar from any *nt* sequence (e-value < 10^-5^, >70% query coverage) or had no hits, these were designated as “Novel PV Types”^32^.

A phylogenetic tree representing the current known diversity of PVs was generated by aligning the amino acid sequences of the 992 known NCBI Virus PV types, and the 383 “Novel PV Types” found in this study over the jelly roll region (Figure 3A). Based on *blastn* searches, 338 PVs were between 60-90% nucleotide identity to *nt* sequences, while 45 sequences had no matches to the *nt* database, likely falling below 50% nucleotide identity. These 45 sequences were clustered at 60% nucleotide, and represent 29 putative new genera (Supplementary File S9).

**Figure 3.**
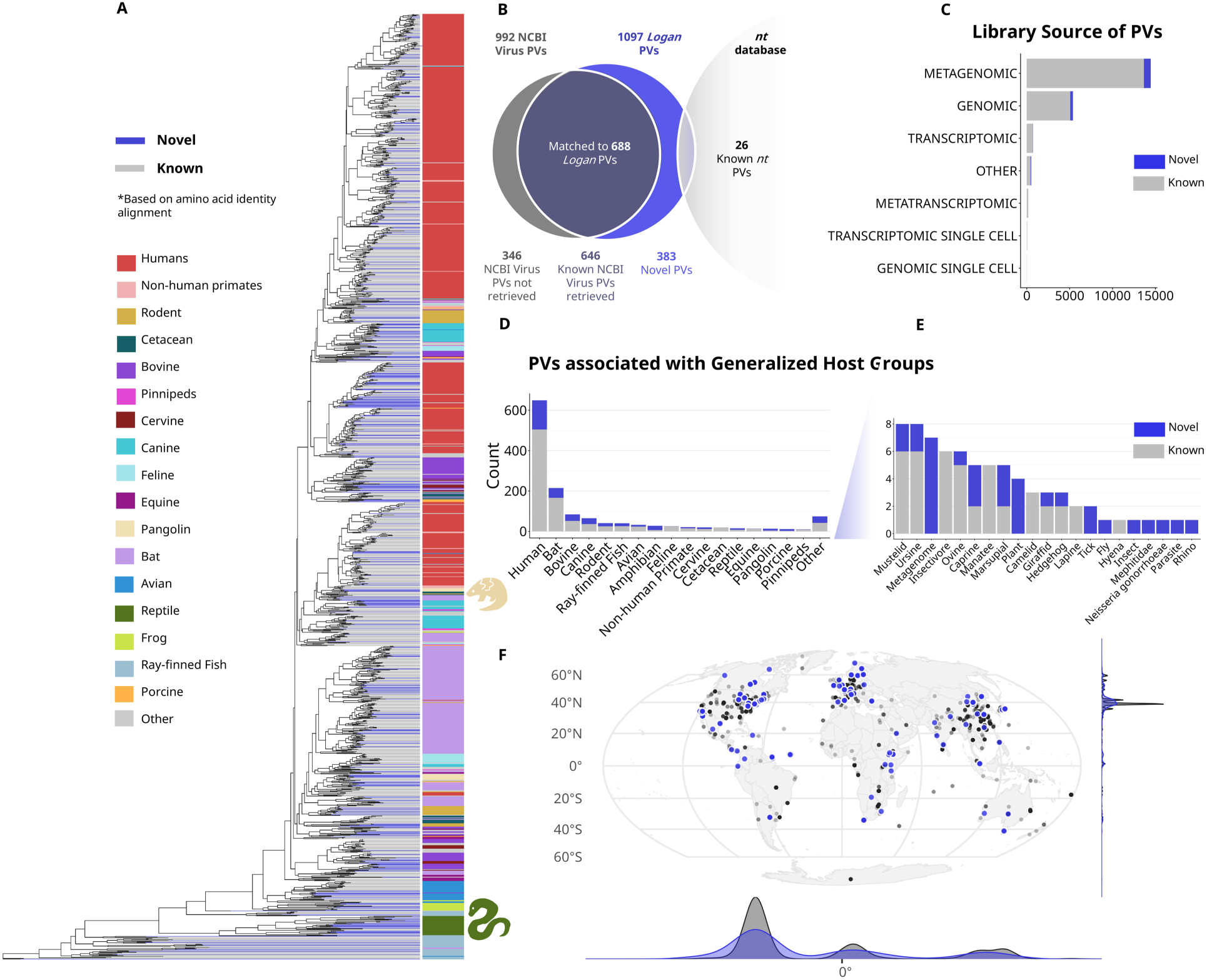
Evolutionary, ecological, and geographic diversity of papillomaviruses. A. Phylogenetic tree of PV L1 jellyroll protein sequences from Logan and NCBI Virus PV types. Sequences were aligned with MUSCLE5 ‘super5’ and the tree inferred with IQ-TREE2 using default parameters, the LG+G4 model, and 10,000 bootstrap iterations^78,79^. The tree was visualized with *ggtree*^80^. Grey branches represent known PV types; blue branches represent novel types identified in this study. Host organism groups (right) were manually assigned from SRA metadata, NCBI or PaVE annotations where available. Organism groups with fewer than nine representatives are classified as ’Other.’ B. Overlap between PV types identified from *Logan* and known PV types. Of 992 types in NCBI Virus, 646 (65%) were recovered by *Logan*. An additional 26 known types were identified through the NCBI nucleotide database (*nt*). A total of 383 novel types were identified. Note that for known NCBI Virus sequences, the known 646 are represented by 688 centroids from *Logan*. C. Library source of *Logan* PV sequences (unclustered). D. Distribution of known (grey) and novel (blue) PV types associated with host organism groups or E. ‘Other’ organisms. F. Geographic distribution of full-length L1+SRA libraries. Coordinates were inferred from BioSample metadata. Marginal histograms show the latitudinal and longitudinal density of detections.

To assess the differences in PV diversity captured within the SRA and NCBI Virus, we compared the types captured in one or both datasets. From *Logan*, 646/992 (65%) of NCBI Virus PV types were independently re-identified, by *blastn* search, using the database consisting of unclustered full length *Logan* L1s (Figure 3B). We further identified 26 PVs only recorded in the *nt* database. This demonstrates that the majority of known PV diversity is present within publicly deposited sequencing data.

In terms of library source, 94% (20,025/21,319) of all PVs in the SRA were found in DNA-based libraries, with 68% (14,460/21,319) originating from metagenomic samples (Figure 3C). In DNA-based libraries, recovered early and late genes were in roughly equal proportion (52.0% E, 48.0% L by *ka.f*), whereas RNA-based libraries were significantly skewed toward early genes (91.3% E, 8.7% L; χ² = 6.3 × 10⁷, df = 1, p < 2.2 × 10⁻¹⁶) (Supplementary Figure S3).

The number of unique known and novel types for each broad host group was tallied, with most types originating from libraries tagged as human samples, followed by bats, bovines, and canines (Figure 3D, E).

Using geographic metadata, the global distribution of known and novel PV types were plotted, with most types being sequenced in major metropolitan areas in the Northern hemisphere (Figure 3F). All underlying data in Figure 3 is available in Supplementary File S7 and S10.

#### Multiple independent observations of metadata can predict virus-host associations

To investigate the biology of the sampled PVs, all full L1s from *Logan* were assigned an associated host taxon. Of the 21,319 L1s, only 735 (3.4%) were not resolved to at least a genus level using the metadata field provided in the SRA library or BioSamples of each sequence, and were manually inspected for contextual information to complete the annotated dataset. Similarly, each of the NCBI Virus sequences were also assigned an associated species, based on NCBI and PaVE annotations. Hosts were grouped broadly into 18 groups at a genus to -class level, reflecting the range of known PV infection (Figure 3). Metagenomes with no defined vertebrate source, organisms not known to be naturally susceptible to PVs such as arthropods or plants, and rare associations (e.g., skunks, shrews) were generalized under the ‘Other’ category. The full table of associations, including original metadata tags and groupings is available in Supplementary File S7.

To assess the reliability of metadata for assigning putative hosts, concordance between SRA-metadata host group associations and NCBI Virus annotations was assessed for the 646 re-identified PV types (Supplementary Figure S4). For PV types that were detected in ten or more SRA libraries or BioProjects, the majority of the metadata labels aligned with the NCBI Virus host, with dense clustering at or close to a concordance proportion of 1 (top right quadrant). Types detected in less than ten libraries or BioProjects showed higher variance in concordance, consistent with lower sample sizes (left quadrants, Supplementary Figure S4A, B).

The majority of PV types on both a per-library basis (n = 598/646, 92.5%) and per-BioProject basis (n = 602/646, 93.1%) were more than 50% concordant with the NCBI-annotated host, demonstrating the applicability of using aggregated, and multiple independent observations of metadata tags to assign putative host groups. Interestingly, a small number of PV types with more than ten observations in both analyses showed concordance below 50% (Supplementary Figure S4A, B).

We manually inspected PVs with NCBI Virus annotated hosts that are discordant with *Logan* metadata annotations (Supplementary Figure S4C). For the porcine annotated Sequence MK377489.1, 90% (89/99) of *Logan* observations are human associated,with no instances of this sequence in the 105,356 porcine samples (all taxid:9821 descendants) within the search space. Conversely, for the human-annotated sequences X70829.1 and MH777222.1, the majority of *Logan* observations were ‘Other’, *Plasmodium* or *Anopheles* samples (n = 68/82, and n = 17/17 respectively). Some sequences such as CS077205.1 and M32305.1 PVs were only found in environmental metagenomic samples, such as soil and cold seep, and host-assignment from complex samples is challenging. Similarly, at a BioProject level, some NCBI Virus PV types appear to have ambiguous or mis-annotated host group associations (Supplementary Figure S4D). The full *Logan* derived metadata for each NCBI Virus accession is available in Supplementary File S8. Together, these observations demonstrate the utility in consensus-based metadata analysis for both assigning putative hosts at scale, and flag misannotations or unexpected host ranges for further scrutiny.

#### The Sequence Read Archive and NCBI Virus capture overlapping but distinct PV types

Of the remaining 346/992 PV types (35%) are not found in *Logan,* 238/346 (69%) had confident matches with e-value < 10^-5^ and >70% query coverage, but were less than the 90% nt-id threshold, representing related but distinct lineages unique to the NCBI Virus dataset.

The additional 108 types (31%) had no confident matches against *Logan* contigs. To determine if certain host groups were disproportionately represented within *Logan* and vice versa, odds ratios (ORs) were calculated comparing the host group distribution of recovered vs. unrecovered types (Supplementary Figure S5). PV types associated with cetaceans (OR = 5.6, p_adj = 0.004), felines (OR = 3.5, p_adj = 0.006), avians (OR = 3.00, p_adj = 0.012), and bats (OR = 1.5, p_adj = 0.0) were enriched among types found only in NCBI Virus. In contrast, amphibians were significantly enriched in *Logan* (OR = 0.29, p_adj = 0.008). The remaining host groups showed no statistically significant difference between those represented in *Logan* or NCBI Virus (Supplementary Table S1). A full description of NCBI-only accessions is available in Supplementary File S11.

#### PVs are present in unique patterns across geographic and ecological space

To understand the ecology of the SRA on a more granular level, the geographic locations of SRA-wide sampling, and of libraries with PVs were summarized and visualized.

Most SRA runs, and consequently PVs, are found in major metropolitan areas, including North America, Western Europe, and East Asia (Figure 4). Based on spatial analysis, the overall distribution of novel PV type proportions is significantly weakly spatially correlated (Moran’s I = 0.06, p < e-5)^33^. Notably, areas statistically enriched in novel PV types compared to overall PV presence are clustered in East Africa and South America, determined by Getis-Ord G_i_* (Figure 4, Supplementary Figure S6)^34,35^.

**Figure 4.**
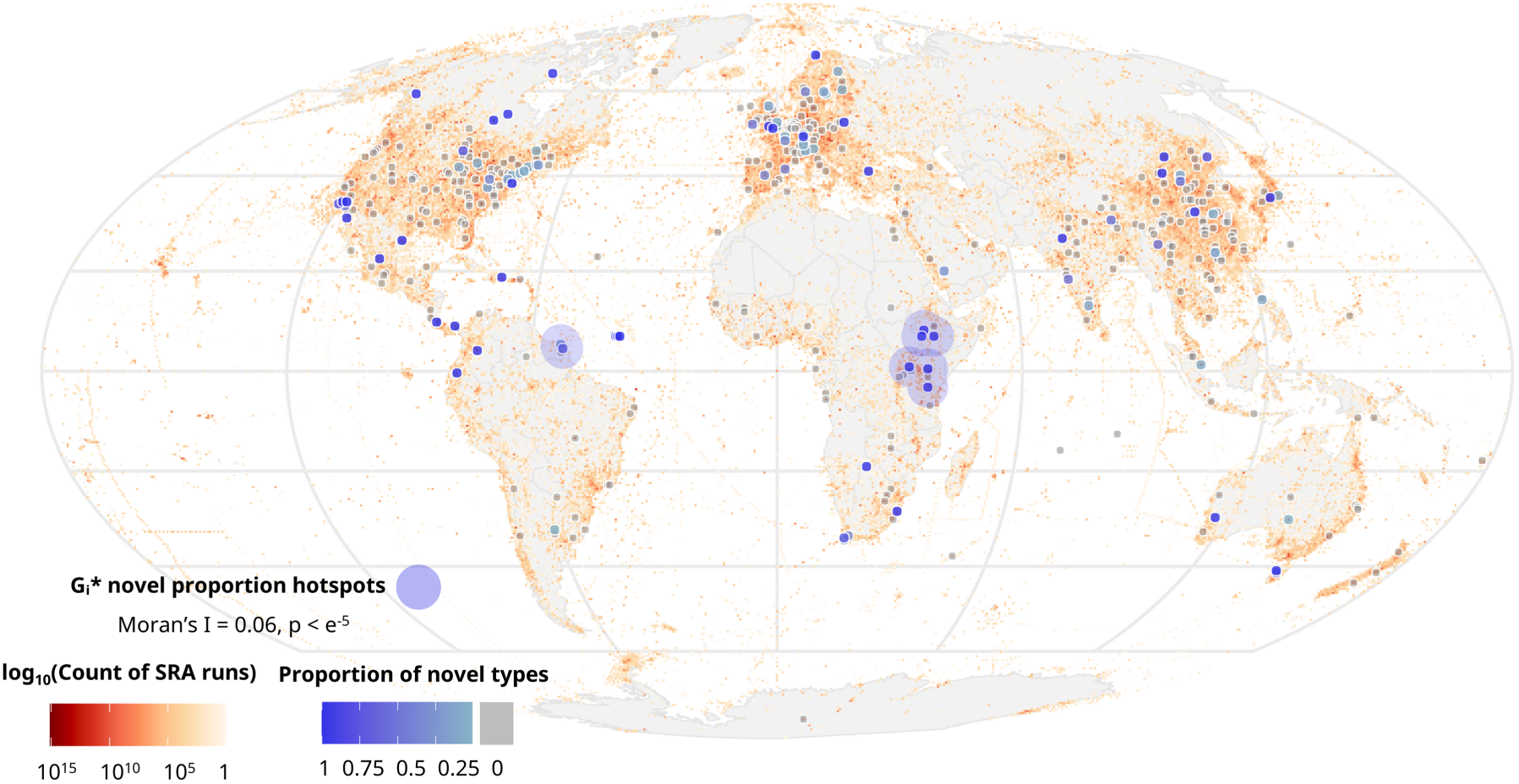
Overlay of PV detections onto SRA sampling density. Each grid square with any novel PVs detected are shaded teal to blue, while those with no novel types are shaded grey. Sampling effort is visualized from white to red, and is concentrated in the regions of North America, Western Europe, and East Asia. Statistically significant hotspots of novel type enrichment identified by Getis-Ord Gi* analysis (blue halos)^34,35^. Spatial autocorrelation of novel type proportion was confirmed by Moran’s I (I = 0.06, p < e⁻⁵)^33^.

In terms of ecological biomes, most sampling efforts in the SRA are concentrated in Temperate Broadleaf and Mixed Forests, accounting for 58% (12,447,825/21,292,703) of all SRA accessions with associated geographic data, consistent with major metropolitan regions (Figure 5A, B). Oceanic sampling follows at 13% (2,826,339/21,292,703), then Temperate Grasslands, Savannas and Shrublands at 12% (2,586,639/21,292,703).

**Figure 5.**
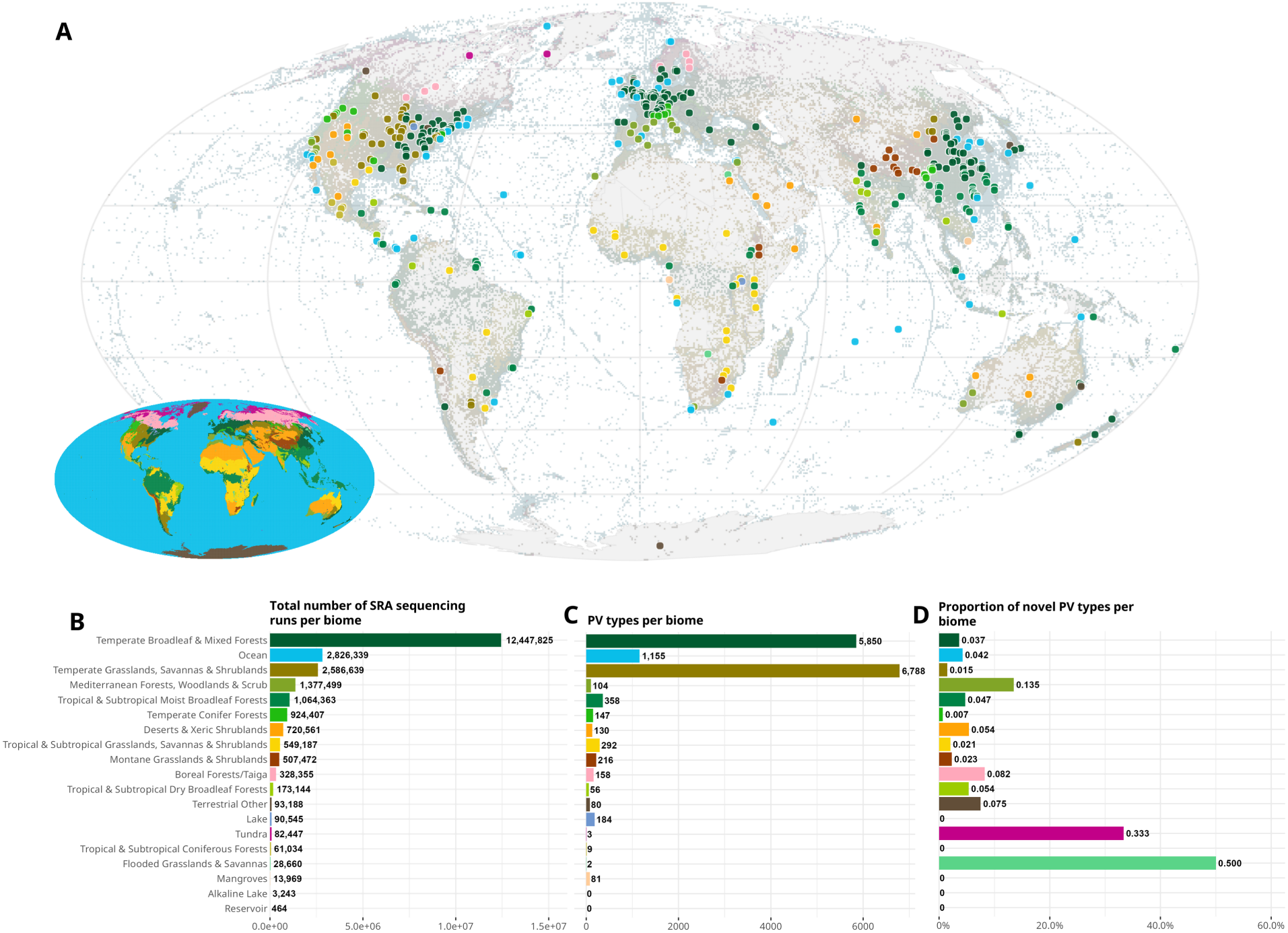
Biome-level distribution of SRA sampling effort and PV discovery. A. Geographic distribution of PV detections coloured by assigned biome. Each buffered grid square contains at least one library with a full-length L1 sequence, underlaid with the biomes of overall sampling efforts in the SRA. Biomes for each grid square were assigned based on the inferred geographic coordinates of the BioSample and geographical overlap with the WWF 2017 Ecoregions. The inset map shows the full global distribution of terrestrial and aquatic biomes used for classification. **B. Total number of SRA sequencing runs per biome, ordered by sampling effort.** Temperate Broadleaf and Mixed Forests account for the largest share of sequencing effort, followed by Ocean and Temperate Grasslands, Savannas and Shrublands. **C. Total number of PV sequences detected per biome, ordered by sampling effort.** Biomes are ordered identically in panels B and C to facilitate direct comparison. **D.** Proportion of novel PV types per biome, calculated as novel/(novel + known).

Most PVs (44%, 6,788/15,613) were sequenced in Temperate Grasslands, Savannas and Shrublands, followed by (37%, 5,850/15,613) in Temperate Broadleaf and Mixed Forest associated accessions (Figure 5C). Relative to sampling effort, more PV types are found in Temperate and Tropical Grasslands/Forests, as well as Lake and Mangrove biomes. Relative to sampling effort, sampling within the Ocean biome produces fewer PV types, while Temperate and Tropical Grasslands, Tropical Forests, lakes, and Mangroves are enriched for PVs. The highest proportion of novel PVs per biome was found in Flooded Grasslands, followed by Tundra and Mediterranean Forest (Figure 5D).

#### Papillomaviruses in Logan and their Molecular, Evolutionary, and Ecological Context

To demonstrate the additional information contained within each PV hit, five representative sequences spanning the tree were manually selected for detailed analysis. Of the five, two were visualized to support the conclusions. The full nucleotide sequence of each contig is presented, along with HMM- (InterProScan hit) or BLAST-identified (e < 10^-5^) ORFs of core and accessory genes^32,36^. Full hit information for each contig is available in Supplementary File S12. The phylogenetic information of each example was extracted, and neighbouring clades are labelled with the associated organism based on manual annotation. Geographic and other metadata were manually curated based on source publications, and every putative L1 gene was folded and aligned to the HPV 16 L1 (PDB: 1DZL) structure in PyMOL^37,38^.

### Predicted gene content of PV case studies

#### Pangolin Associated PV

This SRA accession (SRR25256564) originated from a pangolin re-sequencing project, which extracted genomic material from tissue attached to pangolin scales confiscated by the Yunnan Provincial Forest Public Security Bureau from 2015 and 2019^39^. This sample was identified to be from a white-bellied pangolin (*Phataginus tricuspis*) The Logan-assembled contig (SRR25256564_207500) contains the ORFs for the E1, L1, E2, L2, and E6 genes, and is 5956 nucleotides in length (Figure 6A, Figure 7A).

**Figure 6.**
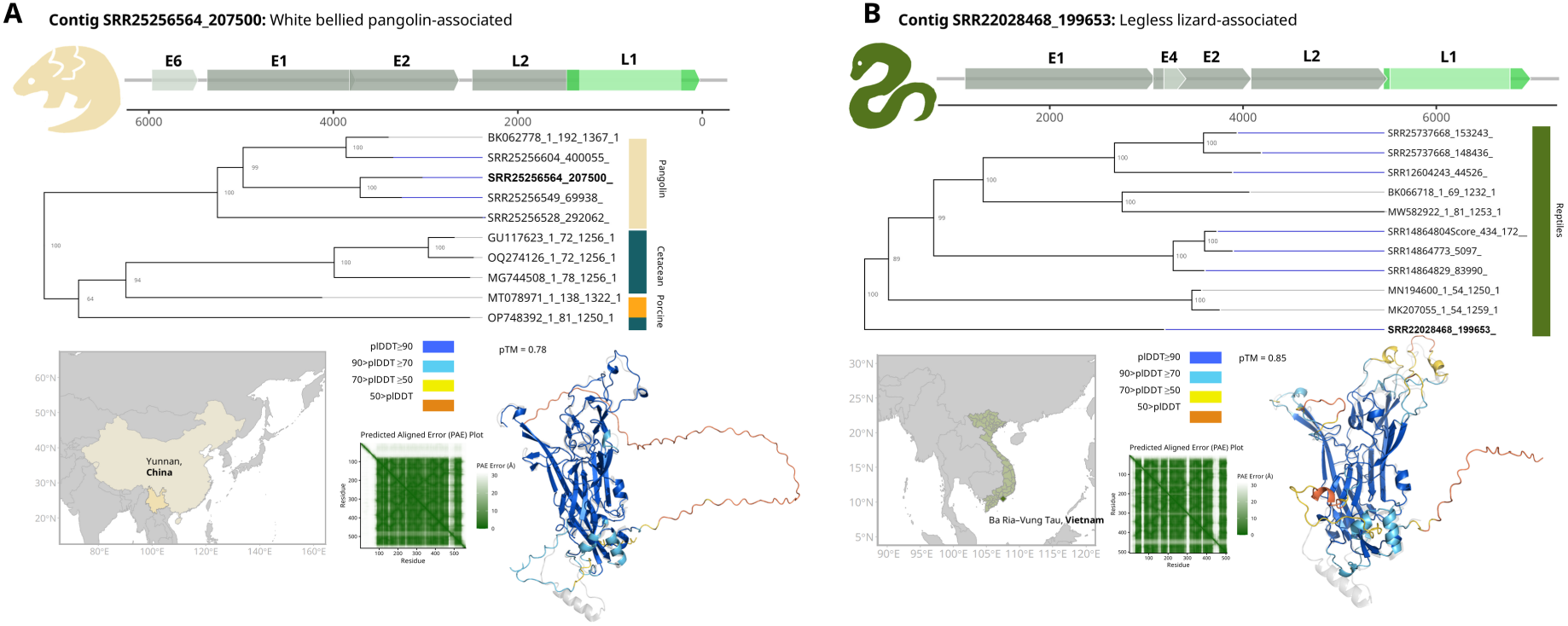
Case studies of novel papillomaviruses from *Logan*. Two representative examples illustrating the biological depth recoverable from individual Logan contigs. **A. White bellied pangolin-associated PV (contig SRR25256564_207500). B. Legless lizard-associated PV (contig SRR22028468_199653).** For each case study, panels show (top) the genomic organization of the assembled contig with identified open reading frames, coloured as in Figure 1A; (middle) a phylogenetic subtree of closely related PV types with host associations indicated by colour, with grey numbers represent bootstrap support after 10,000 iterations, the inferred geographic origin of the source library (bottom left) and (bottom right) the predicted L1 protein structure generated by AlphaFold3, coloured by pLDDT confidence score^88^. The animal icons adjacent to each genome map correspond to the position markers on the full phylogenetic tree in Figure 3A.

**Figure 7.**
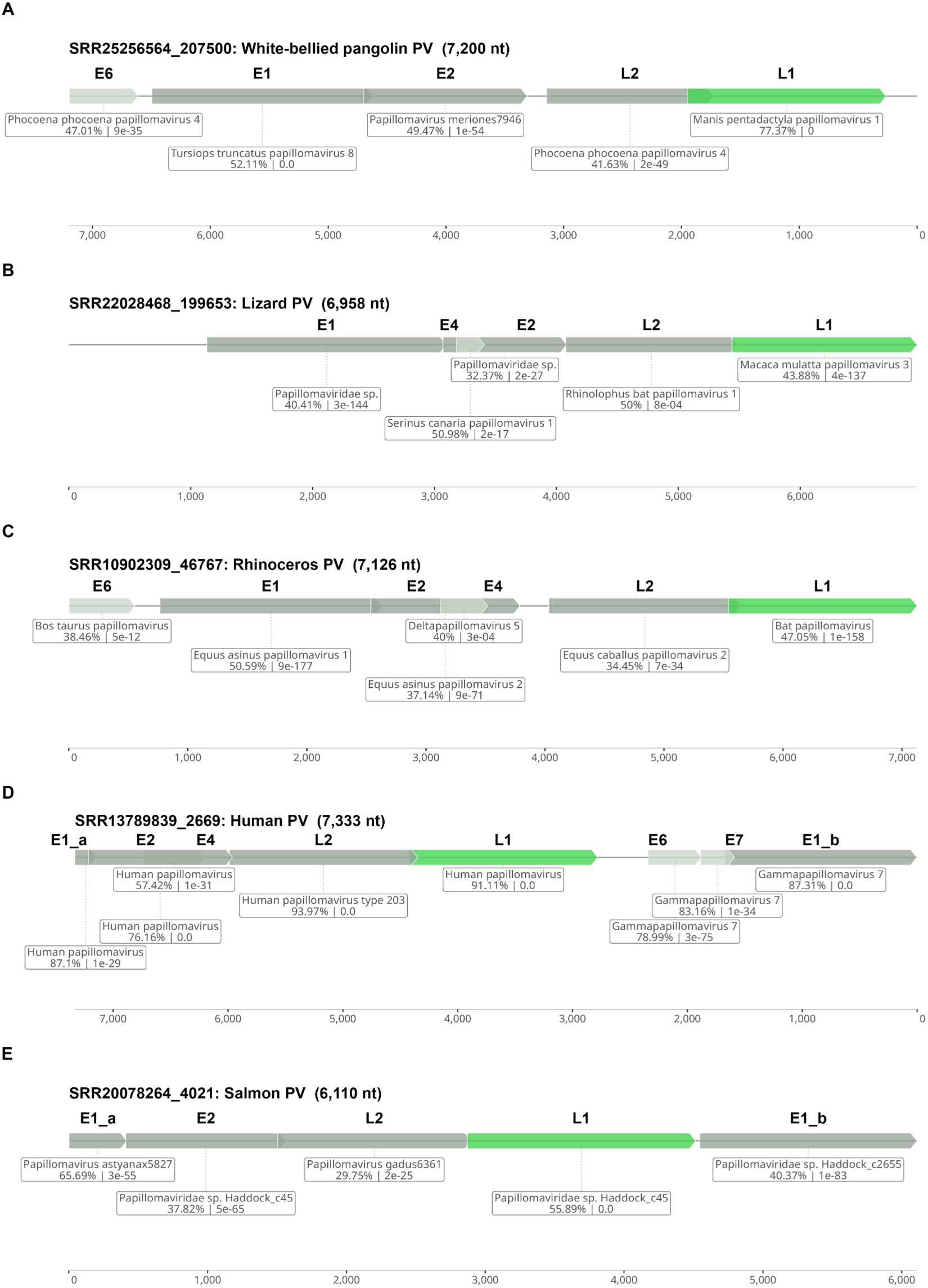
Predicted gene content of five PV case studies. For each PV, stop-stop ORFs were identified by NCBI ORFfinder, and by *blastp* against the *nr* database ^32,107^. For each gene, the top hit is shown with corresponding accession, e-value and amino acid percent identity (%ID). ORFs are visualized based on nucleotide coordinates of the assembled contig.

The L1 amino acid sequence is closely related to a Pathracer-constructed pangolin-associated sequence from the same SRA library, and other sequences from other runs in the same study (SRR25256604), as well as a previously identified pangolin-associated PV from a related, but distinct pangolin sequencing effort (BK062778.1)^40^. The most closely related sister clade contains four cetacean-associated PVs.

The structure of the L1 protein was predicted with high confidence (pTM = 0.78) in the core jelly roll fold and overall aligned closely to the structure of the HPV 16 L1 protein (Figure 6A).

#### Legless Lizard Associated PV

The SRA accession SRR22028468 is linked to a study investigating the evolutionary history of the blind legless lizard *Dibamus deharvengi*, collected from Binh Châu forest in Vietnam, the only habitat in which it is found^41^. For this run, muscle tissue from a female lizard was isolated and sequenced. Only the core genes were identified in contig SRR22028468_199653, which is 6959 nucleotides in length (Figure 6B, Figure 7B).

This sequence represents an expansion of lizard-associated PVs from both NCBI and other independent SRA runs. The predicted protein structure of the core fold aligns closely to that of HPV 16 L1 (Figure 6B).

#### White Rhinoceros Associated PV

SRR10902309 originates from a thesis study aimed at developing genetic tools to reduce inbreeding in southern white rhinoceros^42^. The sample was collected from the skin of *Ceratotherium simum simum* in Botswana – the precise location is obfuscated to protect these vulnerable animal populations. Samples were collected during routine marking and health checks between 1993 and 2013. Contig SRR10902309_46767 is 7588 nucleotides long, and contains 6 identified PV genes, consisting of the core genes and accessory genes E4 and E6 (Figure 7C).

This rhinoceros-associated L1 sequence is placed closely to horse PVs and is part of a sister clade to bat-associated PVs. The Alphafold3 predicted structure of the L1 protein is high confidence, and the core fold closely resembles that of the HPV 16 L1 (Supplementary Figure S7C).

#### Human Associated PV

As part of a study on the impacts of aging on the microbiome, the sample associated with SRR13789839 was collected by swab from the torso of an 87-year-old woman with hypothyroidism and hyperlipidemia in Connecticut, USA^43^. All core genes, including accessory genes E4, E6, and E7 were identified in contig SRR13789839_2669, with a total length of 7335 nucleotides (Figure 7D).

This novel PV type was placed within the Gammapapillomavirus genus, and the predicted protein structure follows that of HPV 16 L1 (Supplementary Figure S7A, E).

#### Steelhead Trout Associated PV

The steelhead trout sample in this study was collected as part of an effort to monitor evolutionary processes within the Columbia River^44^. Specifically, this run is from inland specimens collected within the Rapid River basin. The core genes E1, L1 E2 and L2 were identified in the 6112 nucleotide contig SRR20078264_4021, with the E1 overlapping both ends of the contig (Figure 7E, Supplementary Figure S7F).

Collectively, these case studies represent the information contained within each *Logan* derived contig. In conjunction with the molecular features within the sequence itself, additional metadata such as the phylogenetic context, structural predictions, geographical patterns, and the ecological circumstances can be connected each hit.

## DISCUSSION

### The Sequence Read Archive and Logan as a Frontier of New Biology

Here, we demonstrate that the SRA contains unrecognized PV diversity. We recover two-thirds of known PV types in a single unbiased survey, and record the equivalent of an additional third of novel PV types from 105 putatively associated host species. This work offers a different perspective on the SRA using *Logan*, framing the corpus of public sequencing data not merely as a passive archive but as a vast ecological survey sampling over two-decades of “Earth’s Genome”.

#### The limits of petabase-scale assemblages

While we recovered 65% (n = 646) of known PVs, full length L1s from the remaining 35% (n = 346) were not re-identified in the *Logan* database, and those originate from bat, human and bovine samples. Many of these missed sequences are a result of focused research efforts in wildlife or clinical settings, often using techniques optimized for low-abundance and low-sensitivity detection^29,45–47^. In comparison, public sequencing repositories like the SRA capture an evolutionary breadth of viruses from unexpected taxa. As such, focused and unbiased virus discovery efforts should be viewed as complementary strategies which complement deep characterization of societally- and clinically relevant “known unknowns” with the broad discovery of “unknown unknowns” across evolution.

The choice of using a contiguous sequence of L1-length (∼1000 nt) for taxonomic assignment of PV types is based on taxonomic precedent^22,26^. Requiring a full-length L1 sequence to classify as a PV type caused the most substantial drop in the represented PV contigs, with 4.5% of L1-containing contigs spanning the jellyroll region. For short-read sequencing and assembly such as the major fraction of *Logan-SRA*, fragmentation remains a substantial issue since it requires sufficient read depth to continuously assemble a complete-gene without gaps, but not so much coverage that strain- or sequencing-error introduces branches in the assembly graph^48^. While we show that the graph-aware aligner *PathRacer* can rescue such fragments, this remains computationally costly to perform scale, warranting further development of graph-genome alignment methods.

#### Metadata: Power in (the) Numbers

The *in-silico* assignment of viral hosts is an open challenge in the field^12^. Formal host assignment requires proof of infection, often molecular, while computationally identified sequences rely on sparse and error-prone metadata associations. While the confidence of host-association for the many viruses observed a single time is low; repeated and consistent associations across independent experiments is able to recapitulate, and in some cases improve, curated associations.

92.5% of PV host assignments in *Logan* were concordant with the curated NCBI Virus with a simple 10 observation majority rule classifier. A small number of well-sampled PV types showed unexpectedly low concordance (<50%). Inspection of these cases revealed distinct explanations. The NCBI-annotated *Sus scrofa* PV MK377489.1 appears 89 times in human libraries, and 10 metagenomes within *Logan*, suggesting that this sample may be mis-annotated within NCBI. In contrast, bovine PV AB626705.1, appears 23 times in horse, pig, and rodent *Logan* libraries. While some bovine PVs are known to cause equine sarcoids, the association with porcine and rodent samples may be a result of technical errors (e.g. sample cross-contamination), or previously unknown biological connections^49^.

Human PVs X70829.1 and MH777222.1, are predominantly found in *Plasmodium* (n = 31, 17 respectively) or *Anopheles* (n = 51, 0) libraries, reflecting a unique pattern not observed in other human PVs, though this spans only a single BioProject for both organisms (PRJEB2136 and PRJEB2141). A conclusive association cannot be drawn here due to the lack of independent replication, but this raises the question if some PVs can persist or even spread within mosquito vectors. Some evidence suggests that mosquitoes and ticks can mechanically transmit rabbit PVs after feeding on an infected host, but this mode of transmission has not yet been demonstrated for human PVs^50^.

Finally, of interest are the PV types with multiple observations, yet lower concordance relative to other types within the same range of observations (Supplementary Figure S3A, B), which may represent PVs with host ranges that span multiple host groups, warranting further scrutiny.

Collectively, these examples suggest that using existing metadata over independent observations of a PV type provides a means to validate putative host associations, and also a method for flagging potential misannotations. As sequencing data is expected to grow exponentially and the number of discovered viruses expands, computational assignment of host-ranges through sequence meta-data may prove to be an effective strategy to predict virus host-ranges, especially when combined with phylogenetics.

### The Sequence Read Archive Expands Known PV Diversity

The phylogenetic placement of most novel PV types from *Logan* represents expansions to known clades, rather than new lineages. The clade host-associations we observe are consistent with the model of the evolutionary origins of PVs arising 424 million years ago, around the Devonian period and parallel the diversification of fishes^15^. What the distribution of novel PV types suggests is that existing PV diversity is unevenly sampled rather than fundamentally unexplored. NCBI Virus-described PVs are enriched in bat, cetacean, feline and avian PVs, likely reflecting anthropogenic study bias and the disproportionate viral surveillance programs in these species [53, 54,55,56,57].

In contrast, *Logan* PVs are enriched for amphibians, as they largely lack perceived zoonotic or industrial relevance and are not a focal group for viral discovery relative to mammals and aves (Supplementary Figure S5)^51^. Notably amongst the other 105 putative host-associations for novel PV types were species such as the southern white rhinoceros, voles, and grey foxes^27^. Interestingly, most highly-diverged PVs were found in reptilian and amphibian-associated libraries, suggesting these hosts represent disproportionately more PV diversity in comparison to well-sampled taxa (Supplementary Figure S8).

While such metadata-derived host associations should be interpreted cautiously, improved characterization of the wildlife virome can support conservation efforts for endangered and at-risk species, especially given the potential oncogenic effect of known PVs in human and non-human species. The range of potential host diversity captured here highlights its capacity for systematic discovery to uncover ecological relationships.

Beyond host associations, the vast majority of PVs (94%, n = 20,025/21,319) in the SRA were identified from DNA-based libraries, with metagenomic samples representing the largest single source (68%, n = 14,460/21,319). First, this highlights the utility of ‘untargeted’ metagenomic studies for viral discovery. While the original intentions of these studies may not be focused on sequencing viruses, their collective effectively forms a passive virome surveillance network which can be interrogated through *Logan*.

Second, the enrichment of PVs in DNA libraries likely reflects the methodological limitation of L1 as the hallmark PV gene. In PVs, the late (L) structural genes L1 and L2 are primarily expressed during short periods of productive viral replication and are not constitutively transcribed during early infections, or the long latency periods of PVs, unlike the early (E) genes^18,52^. As a result, L1 sequences in the SRA are more likely to be captured as DNA, either from virions co-sequenced with tissue or integrations into host cell genomes. This is consistent with the observation of low representation of L1 sequences relative to E1 in RNA-based libraries (Supplemental Figure S3). If PV taxonomy was performed on an early gene expressed throughout the viral lifecycle, such as the conserved E1 helicase, this would recover additional PVs more putative hosts, in which virion production is inactive, but persist latent^53,54^. Future sequence mining can be optimized through the consideration of genomic versus transcriptomic representations of targets.

#### Papillomaviruses as case studies

The case studies of individual PV hits presented here illustrate both the breadth and the depth of information recoverable from a single *Logan-*derived contig, and the diversity of contexts in which viruses can be incidentally captured. From these examples, not only are full L1 genes retrieved, but also near- or fully complete PV genomes.

#### Pangolin-associated PV

In the case of the PV associated with the endangered, white-bellied pangolin, the sequencing library was generated from the tissue attached to a confiscated pangolin scale in China, for the purposes of pangolin genome re-sequencing^55^. A contig containing five identifiable PV genes, including the L1 hallmark, was identified through *Logan* search. The predicted structure of the L1 protein closely aligns with that of the HPV 16 L1, highlighting the high conservation of this protein throughout all PVs. The L1-based phylogenetic placement of this pangolin-associated PV groups it closely with another pangolin-associated PV (BK062778) that was independently identified through re-assembly of a separate SRA library (SRR9018603) focused on pangolin genomes, as well as another pangolin-associated PV recovered from *Logan*^56,57^. The sister clade to these pangolin PVs consists of cetacean- and wild boar-associated types. More broadly, this pangolin-cetacean-boar group falls within a larger clade dominated by human Alphapapillomaviruses.

Interestingly, this is not the only place on the phylogenetic tree where pangolin-associated PVs appear. A second group of pangolin-associated PVs are positioned basally, adjacent to shrew-, bat-, and rodent-associated PVs. This group consists of PVs from both SRR900505[3-6] libraries, and SRR25256[542, 569, 617] libraries, which were found to originate from separate studies carried out by the same group in Yunnan^39,58^. The placement of pangolin PVs in two distant clades lends support to two possible interpretations. This could reflect biologically distinct PV lineages that have independently adapted to pangolins as hosts, a pattern that has been well established in cattle, where ∼4 phylogenetically distant PV genera (Delta-, Xi-, Epsilon-, and Dyoxipapillomavirus) have been found to circulate within the same host species^46^. Alternatively, the phylogenetic placement may indicate technical artifacts. Both pangolin sequencing efforts originate from the same research group with partially overlapping authorships, albeit linking to separate BioProjects, submission dates, and publications 3-4 years apart. This raises the possibility that cross-contamination during sample processing, laboratory work, or sequencing libraries could account for the discrepancy. It is also possible, though perhaps more unlikely, that biological material between cetaceans and pangolins, which are trafficked through similar routes and networks, may have come into physical contact before laboratory acquisition. Resolving this ambiguity requires independent sampling from confirmed pangolin tissue under controlled conditions, but the observation itself underscores the biological insights that can be derived from SRA-wide analyses and provides an avenue of further study.

#### Legless lizard-associated PV

The PV associated with legless lizards was recovered from a study aimed at understanding the evolutionary history of sex determination in squamate reptiles, representing an expansion of reptilian PV diversity from one of the most undersampled vertebrate groups. This sequence was assembled from the muscle tissue of a female *Dibamus deharvengi*, from Binh Châu forest, the only known habitat for this species of lizard^59^. The recovery of a PV from this species is noteworthy given the natural history context of this animal, which has never been the subject of virological investigation to the best of our knowledge, and further provides an excellent example of the unpredictable nature of which datasets may provide novel biological insights. Interestingly, the source material for this sequencing run was annotated as muscle tissue, which contradicts the expected tropism of epithelium. Although in some cases human PV DNA has been detected in peripheral blood and other non-epithelial compartments, for small specimens, co-extraction of skin during tissue dissection remains the more parsimonious explanation^60,61^.

From an ecological perspective, this represents the first record of a PV associated with these geographically limited species. While there is no evidence of pathology in this case (the authors did not note any physiological anomalies in the collected lizards), PVs in other vertebrates are known to opportunistically progress from latent states to disease in stressful conditions where the immune function of the host is impacted^62^. Climate change has progressively applied increasing pressure to habitats globally, and this is of particular concern to these range restricted lizards, providing a baseline in which to understand the potential virus-host interactions that may occur in the face of habitat pressure.

In addition, the known diversity of small DNA viruses, including PVs, in “lower vertebrates” such as reptiles and fish have historically been poorly characterized. Only a handful of reptilian PV (∼5) genomes have been recorded to date [2025]^27,63–65^. The lack of representation in this area of the PV phylogeny has impacted the ability to draw conclusions about PV evolutionary history, introducing uncertainties in processes such as adaptive radiation, host jumping, and even the origin of the PV oncogenes E6 and E7^66^. Thus, each new reptilian or “lower vertebrate” PV sequence contributes disproportionately to understanding the trajectory of PV evolution.

### The Sequence Read Archive Reveals the Geographical and Ecological Distribution of PVs

The geospatial distribution of PV sequences from the SRA reflects the global distribution of total sequencing efforts. The highest PV-detection density is concentrated in North America, Western Europe, and East Asia. Perhaps unsurprisingly, these regions make up the majority of SRA depositions and reflect the socio-economic factors that determine where and what is sequenced.

More interestingly, the proportion of novel PV types uncovered in each region does not recapitulate overall sampling density and is not equally distributed globally. Through permutation testing, significantly enriched grids of PV novelty were identified in areas of lower sequencing effort (Supplementary Figure S6). Regions in South America and East Africa were identified as weak, but statistically significant hotspots (Getis-Ord Gi*, detecting spatial clustering of similarly high values rather than isolated outliers), characterized by spatially clustered grid squares with consistently high proportions of novel types, in contrast to isolated points of novelty surrounded by well-sampled grids. This inverse relationship between sampling effort and novel enrichment suggests that the least-sampled regions likely harbour more PV diversity, consistent with the broader concept that continued sampling in well-studied regions yields diminishing returns^67^. This negative correlation provides a basis for prioritizing future efforts in regions with low sequencing representation but high novel ratios, both in terms of fieldwork and computational re-analysis.

By integrating an additional layer of biome information to the geographic locations of SRA samples, we reveal a disconnect between where PVs are discovered and their biomes. Temperate Broadleaf and Mixed Forests accounted for the largest share of SRA libraries, consistent with the major sequencing centres and clinical research institutions situated in the temperate regions of North America, Europe, and Asia. However, for PVs, Temperate Grasslands, Savannas and Shrublands, followed by subtropical and tropical grasslands, freshwater lakes, and mangroves yielded the highest proportion of PV types per unit of sampling effort.

The mismatch between sampling effort and discovery poses and interesting question: is this pattern a result of the biological underpinnings of these biomes, such as host diversity, biomass, and organism density, or is it more likely to be a methodological artefact-driven differences in the types and design of experiments conducted in these regions? For instance, it is possible that libraries from grasslands, tropical, and aquatic ecosystems are more likely to have been collected for the express purposes of ecological monitoring and metagenomic studies, which are likely to incidentally capture PVs. Temperate broadleaf libraries, especially those aligning to major metropolitan regions, are likely dominated by clinical and biomedical research efforts, which may be more targeted and thus biologically ‘narrow’.

Without controlling for host species richness and more granular assignments of samples to biomes, disentangling these relationships is beyond the scope of this study. Nevertheless, the observation that the discovery of PV types is not proportional to sampling efforts across biomes can inform where future PV discovery efforts should be focused.

## CONCLUSION

Using the *Logan* assemblage for the systematic survey of the SRA we recovered 383 novel PV types across 105 associated host species, expanding known PV diversity by 34%. In parallel, 65% of all known PV types were re-identified. This demonstrates how a single 10-hour *Logan* search yields the data to recover the majority of five decades of PV sequencing efforts, and adds to this corpus. This, along with other petabase-scale virus discovery studies [ref serratus], support the concept significant virus discovery advances can be made through cost-effective data-reanalysis.

Importantly, this work extends beyond virus biodiversity discovery and establishes a framework for integrating source organism, geographical, and ecological data across the SRA. By integrating this information, we were able to infer information past that of PVs evolution alone, making predictions about putative hosts, geographic and biome-level distributions. Additional avenues of exploration, such as temporal trends, pathogen and host co-occurrence, and finer-scale ecological modelling are all tractable with the pre-existing information.

More broadly, this framework can be extended to any viral or gene family. The jellyroll fold is shared by PVs, polyomaviruses, and other DNA viruses providing an entry point for describing the total diversity of DNA viruses across the SRA^25^. As *Logan* is updated with new data, and metadata sparsity decreases, the comprehensiveness of these analyses will grow accordingly, providing the data for future virus foundation models.

By systematically accessing the collective sequencing efforts of researchers globally, the challenge is no longer in producing new data, but to develop the infrastructure and methods to extract the rich biological information already present.

## METHODS

### Extracting Known Papillomavirus Sequences from NCBI Virus

To establish a baseline of known PVs within NCBI, all PV nucleotide sequences (*Papillomaviridae*, taxid:151340, as of 2024-06-18) in the NCBI Virus database were downloaded (n = 44,909; nucleotide PV sequences) ^68^. Associated metadata for each sequence was included in sequence headers using the NCBI Virus ’Species’ and ’Organism Name’ options during download. EMBOSS v6.6.0.0 *getorf* with default parameters was used to retrieve open reading frames (ORFs) > 30 nucleotides long, translating sequences between stop codons (n = 5,067,231; stop-stop ORFs)^69^.

To confirm that each nucleotide sequence corresponds to a known PV protein, ORFs were annotated using 11 PV Pfam profile Hidden Markov models (HMMs) (Table 1) and HMMER (v3.4) *hmmsearch* with options ‘--F1 0.01 -T 20 --domtblout’ to confirm they were of PV origin^70,71^. ORFs with bitscore > 20 to core gene (E1, E2, L1, L2) HMMs (n = 57,107; HMM-filtered ORFs) were kept for downstream processing.

**Table 1.** HMMs retrieved from Pfam for annotation of PV sequences from NCBI Virus.

| Pfam Entry | Name | PV Gene | ORFs from NCBI Virus |
| --- | --- | --- | --- |
| PF00524 | PPV_E1_N | E1, N terminal domain | 11,306 |
| PF20450 | PPV_E1_DBD | E1, DNA-binding domain | 11,670 |
| PF00519 | PPV_E1_C | E1, C terminal domain | 11,891 |
| PF00508 | PPV_E2_N | E2, N terminal domain | 11,295 |
| PF00511 | PPV_E2_C | E2, C terminal domain | 11,875 |
| PF00500 | Late_protein_L1 | L1, major capsid protein | 21,800 |
| PF00513 | Late_protein_L2 | L2, minor capsid protein | 11,798 |
| PF02711 | Pap_E4 | E4, early protein | 9,842 |
| PF03025 | Papilloma_E5 | E5, early protein | 9,195 |
| PF00518 | E6 | E6, early protein | 17,351 |
| PF00527 | E7 | E7, early protein | 14,665 |

To reduce computational redundancy in petabase-scale search, these ORFs were clustered at 90% amino acid identity (n = 4,207) with *usearch* (v.11.0.667_i86linux32) ‘-cluster_fast -id 0.9’.

### Trimming of non-viral flanking sequences

PVs are double-stranded DNA viruses that replicate as episomes within the host cell nuclei, and can occasionally integrate into the host genome if the viral lifecycle is dysregulated^72,73^. Consequently, PV nucleotide sequences may contain flanking host-derived sequences adjacent to viral ORFs.

To remove this source of non-PV contamination (flanking genomic integration sites, cloning vectors, and insertions) in PV ORFs a translated nucleotide search with DIAMOND *blastx* (v2.1.8.162) against the human (*hs1*), mouse (*GRCm39*), cow (*bosTau9*), yeast (*sacCer3*), and E. coli (*GCF 000005845.2 ASM584v2*) reference genomes was carried out. Briefly, contigs of 1000 nucleotide sliding windows with 200 nucleotide overlaps of each reference genome above were generated with *seqkit* (v2.8.0) ‘sliding -s 800 -W 1000’ to simulate a database of common sequence library contaminants^74^. These genomes do not comprehensively screen against all non-viral sequences, but represent the most common false positive ‘contaminants’ during initial pilot testing. The decontamination alignment was completed using DIAMOND *blastx* with parameters ‘--masking 0 --ultra-sensitive --seed-cut 0.9 -p 8 -k1 -f 6 qseqid qstart qend qlen qstrand sseqid sstart sendslen pident evalue cigar qseq translated full qseq full qseq matè^75^.

### Creating PVDB1, a protein sequence set for petabase-scale sequence search

All amino acid-clustered PV ORFs (n = 4,207) were screened for regions matching non-viral genomic sequences with e-value < 10^-10^. Genomic matches were trimmed from ORFs, by manually truncating from either the start or end of the sequence, depending on the location of the contaminating region (n = 9), with bedtools (v2.31.0)^76^. ORFs with less than 50 amino acids remaining after trimming were removed (n = 7). Coordinates for trimmed ORFs from NCBI Virus are supplied in Supplementary File S2. This yielded a final 4,200 HMM-trimmed and clustered ORFs for our search query dataset, PVDB1.

### Creating a PV nucleotide sequence set for L1 taxonomic categorization

Using the HMM-filtered NCBI Virus PV dataset, the full jellyroll region of L1 sequences were extracted through using a set of custom HMMs for the substructures of the L1 protein. These custom HMMs (jrHMM) were based on the jellyroll regions of the NCBI PV Virus and PaVE sequences, clustered at 90% amino acid similarity to reduce the over-representation of human PVs. This custom jrHMM is available in Supplementary File S3. The resulting 11,055 L1 sequences were clustered at 90% <u>nucleotide</u> identity with *usearch* (v.11.0.667_i86linux32) ‘-cluster_fast -id 0.9’, a species-like demarcation used in the literature to define the known PV ‘type’, this resulted in a total of 992 known PV types. This clustering threshold is consistent with the criterion used by the International Committee on Taxonomy of Viruses and the Papillomavirus Episteme (PaVE) for defining PV types. This ‘type’ taxonomic level is consistent with clinically familiar designations, such as HPV 16 or HPV 18. Of note, PV can be further grouped into ‘variants’ at 98% nucleotide identity^27^.

### Search Iteration 1: PVDB1 search in the Logan v1.0 assemblage

Using PVDB1, DIAMOND *blastx* with parameters ‘-c 1 --masking 0 --target-indexed –sensitive -s 1 --evalue 1e-8 -k1 -f 6 qseqid qstart qend qlen qstrand sseqid sstart send slen pident evalue cigar qseq translated full qseq’ was run against the the *Logan* v1.0 contig assemblage [2024-07-13, 26.8 million accessions, 385 terabytes of contigs], resulting in 1,931,672 contigs matching input sequences (e-value < 10^-10^)^10,75^. Following the methods above for ORF identification and annotation with HMMs, 26,283,764 stop-stop ORFs culminated to 1,168,298 putative PV ORFs (Supplementary Figure S1, Supplementary File S4).

### Search Iteration 2: PVDB2 search in the Logan v1.1 assemblage

To increase the sensitivity of the search, PVDB1 and all core genes identified from *Logan* with PVDB1 with more than 50 amino acids were clustered at 90% amino acid identity (n = 29,156 centroids from *Logan* v1.0, n = 1,582 centroids from NCBI, n = 30,738 sequences total). This combined dataset (PVDB2) was then similarly decontaminated as above (n = 28 trimmed, n = 33 removed in Supplementary File S1). In parallel, the *Logan* assemblage was updated to increase contiguity (v1.1).

Using PVDB2 (n = 30,705), the *DIAMOND blastp* search was rerun (as above) on the Logan v1.1 database [2025-02-04, 315 terabytes of contigs], resulting in 2,763,517 PV-matching contigs (e-value < 10-^-10^) and 1,365,855 HMM-filtered PV ORFs (Supplementary File S5).

### Characterizing the Evolutionary Landscape of Papillomaviruses

Full length L1s (defined as the start of the B sheet to the end of the I sheet of the jellyroll fold) were identified by jrHMM as above. From 307,441 contigs annotated as containing an L1 ORF (11% from all hits), 4.5% (14,048) contained all eight beta sheets in the full jellyroll structure (Figure 2).

To saturate the L1 search, a custom build of the SPAdes assembler (branch https://github.com/ablab/spades/tree/graph-extract) and PathRacer (included in SPAdes v4.0.0) was used to extract subgraphs and close technical gaps^31,77^. To avoid highly complex and computationally costly searches, PCR amplicon libraries were removed, filtering on the ‘Source’ and ‘Strategy’ metadata columns for each SRA library (n = 1,375/45,258 removed). These libraries are likely to have many single nucleotide polymorphisms or errors caused by the amplification process and are not computationally tractable to search at scale with graph traversal.

Briefly, utilizing graph linkage information in the headers of *Logan* derived contigs, the underlying *de Brujin* assembly graphs for libraries with >= 3 hits (n = 18,642) to individual jrHMM regions (6 total regions, B, CD, E, F, GH, and I) were reconstructed, representing cases where a full, but fragmented L1 may be present. To ‘traverse’ these graphs, PathRacer identifies connections between the ends of contigs that when combined, correspond to full-length L1s, defined by HMM. This process can recover the fragmented contigs from the original library that were not successfully resolved. This fragmentation can occur during assembly as a result of biological variation (e.g., single nucleotide polymorphisms, branching due to variable regions, low virus copy number), or technical limitations (simplified assembly pipelines utilized by *Logan*). Paths that resulted in sequences with > 95% coverage of the full jrHMM, run with command ‘logan-extract -d 1000 -t 8’ and ‘pathracer --length 0.95 --top 5 --threads 16 --rescorè, were retrieved, resulting in an additional 7,275 full L1 sequences.

The nucleotide sequence of the jellyroll region, based on jrHMM-identified coordinates, or directly from PathRacer, was extracted from all full L1s from *Logan* (n = 21,323) to standardize contigs for similarity searches. Upon inspection, four sequences originating from the library SRR6976994 were removed, due to reconstructed contigs containing many simple repeats (n = 21,319 after removal). These sequences were then clustered at 90% nucleotide identity, resulting in 1,097 sequences representative of PV types in *Logan* (centroids) (Figure 2).

These centroids (n = 1,097) were then searched against the *nt* database using *blastn* with options ‘-outfmt “6 qseqid qstart qend qlen qstrand sseqid sstart send slen pident evalue” -evalue 0.001’ [*nt* database as of 2025-02-10]^32^. Hits with less than 70% query coverage between full *Logan* L1s and NCBI sequences were removed, based on the observed bimodal distribution separating partial from full-length alignments, to avoid false negatives arising from alignments consisting solely of short, highly conserved regions (Supplementary Figure S2).

Classification of novelty is as follows: L1 sequences with more than 70% query coverage, **a)** high confidence matches to known PVs (e-value < 10^-5^) but less than 90% nucleotide identity, or **b)** had no significant matches (e-value > 10^-5^) to any sequence in the *nt* database, were considered novel. Comparisons against the NCBI Virus types were carried out by generating a nucleotide database consisting of all full L1 sequences from *Logan* and searching using *blastn* as above. The same filtering parameters (> 70% query coverage, e-value < 10^-5^ and < 90% nucleotide identity cutoff) were used to determine if NCBI Virus PV types were re-identified. Sequences with no hits (n = 163) using default *blastn* parameters were re-searched using ‘-task blastn’ to identify matches with more distant homology. After re-searching, sequences with no match (n = 45) were considered to be at least < 50% nucleotide identity to known PVs. For each novel PV, the closest nucleotide identity match at 70% coverage of the L1 gene was considered the best representative in the *nt* database.

To construct an informative phylogeny, all novel L1 sequences from *Logan* (n = 383, at 90% nucleotide identity), plus all known types from NCBI Virus (n = 992, at 90% nucleotide identity) were translated to amino acids. Some sequences (n =11) were identical at the amino acid level within the jelly roll region. A total of 1,375 unique sequences were retained for the tree. All sequences were aligned with MUSCLE5 (v5.1 Linux) with option ‘-super5’^78^.

IQ-TREE2 (v2.4.0 Linux) was used to generate a 10,000-bootstrap phylogeny with parameters ‘-B 10000 -keep-ident’ and ‘-mset LG,WAG,JTT,HIVb,HIVw,FLU,RTREV -mrate G4,G4+F’ to determine an appropriate model^79^. The resulting tree (using model LG+G4) was visualized and annotated with R packages *ggtree* (v3.12.0) and *tidyverse* (v2.0.0)^80,81^. To refine the metadata associated with each PV, DIAMOND *blastp* ‘--masking 0 --sensitive -s 1 -c1 -k 10 -b 5 --threads 16 -f 6 qseqid qstart qend qlen qstrand sseqid sstart send slen pident evalue full_qseq staxids sscinames’ was used to search each sequence against the *nr* database [updated 2025-04-14] to identify the closest related sequence^75^.

### Annotations and Metadata Crowdsourcing

To assess the reliability of using SRA-derived metadata for inferring host group associations, a concordance analysis was performed with the 646 NCBI Virus PV types re-identified in Logan.

For each known PV, all unique SRA libraries containing a full L1 sequence that aligns to a known PV with more than 90% nucleotide identity, 70% coverage, and e-value < 10^-5^ were identified (n = 8,524). The organism metadata field associated for each SRA library was recorded (n = 6,138). In cases where the host metadata field was not available within the SRA library, the host or organism field linked BioSample is used instead (n = 1,651). Finally, for libraries that have no relevant fields included, or are labelled ambiguously (such as ‘metagenome’, ‘viral metagenome’, ‘skin metagenome’) were manually inspected and curated to reflect the sample origin (n = 735). For instance, a sample labelled only ‘metagenome’ may indicate in sample descriptions or comments that it was collected from a human skin swab, and thus would be re-labelled as originating from a human sample.

To reduce bias introduced from libraries with high sequencing depth or single samples, this ‘concordance’ analysis between the NCBI-annotated host, and metadata labels from *Logan* was calculated on both a per-library, and per-BioProject basis. The proportion of libraries or BioSamples that were labelled with the same generalized host group of the expected host was calculated per PV type. This concordance value was then plotted against the number of observations of the library or BioProject that contains that PV type, using *ggplot2*.

For known PV types, the NCBI records associated with the accession and PaVE annotations, were used together to manually assign an associated organism. For novel PVs found in this study, the ‘organism’ SRA metadata field was used to manually assign associations where possible, descriptions included within the BioSample description, or associated publications, in order of priority.

To characterize the taxonomic novelty of putative PV clusters identified above by host association, we binned the 1,097 centroid clusters by their best L1 nucleotide identity to the *nt* database in 5% increments (50–100%). Centroids lacking a *blastn* hit above the query coverage threshold (≥70% for *megablast*; ≥50% for a secondary search using *blastn* task with word size 11) were assigned to a separate “no hit” category.

For each host group and identity bin cell, we tested for overrepresentation using a two-sided Fisher’s exact test on the 2×2 contingency table (host in bin vs. other hosts; this bin vs. other bins), with Benjamini–Hochberg adjusted p-values. Significance indicators were overlaid on bins where a host was significantly enriched (adjusted p < 0.05, observed/expected > 1).

For interpretability and generalization, each associated host species was manually grouped at roughly between the class and genus levels based on NCBI taxonomy labels (e.g., canine, feline, bovine), with the exception of non-human primates and humans (Supplementary File S6). Organism groups with less than 9 representative PV types (n = 21 groups) were clustered within the ‘Other’ category for visualization.

### Papillomavirus metadata annotation

The library source (transcriptomic, genomic, etc.), and BioSample of origin for each accession was determined using the *efetch* (v20.4) command line program with the SRA run accession ‘-db sra -id “$(paste -sd, sra_runs.list)” -format runinfò^82^.

To assess whether library preparation strategy influences the recovery of papillomavirus genes, we quantified the relative abundance of PV ORFs across SRA library types. The 1,365,855 HMM-filtered PV ORFs were annotated against the Pfam PV gene profile HMMs (Table 1) using HMMER as above. The highest-scoring HMM was used to classify the ORF. Because E1 and E2 are represented by multiple HMM sub-profiles (E1_N, E1_DBD, E1_C and E2_N, E2_C, respectively), the averaged contig-level abundance metric (*ka.f*, a k-mer-based approximation of sequencing coverage reported by Logan^10^) across sub-profiles within each contig to avoid inflating their contribution. Each SRA run accession was classified by library source using metadata retrieved from *efetch* and the ENA Portal API^32,83^. Gene proportions were computed as the sum of *ka.f* values per gene divided by the total *ka.f* within each library type. To test whether DNA-based and RNA-based libraries differed in their recovery of early (E) versus late (L) genes, a chi-squared test was performed on the 2×2 contingency table of aggregated *ka.f* values.

For each SRA-derived PV sequence, refined geographic coordinates inferred from BioSample metadata were assigned to each unclustered full PV L1 hit from *Logan* and Pathracer (n = 21,319 in total). As above, a *blastn* search was conducted on the jellyroll region to determine novelty, with the same classification criteria. As location data can be specified by multiple metadata fields, attribute fields were prioritized in order of accuracy and precision: “lat_lon”, “geographic location (latitude), geographic location (longitude)”, “geo_loc_name_sam”, “geo_loc_name”, “geo_loc_name_country_calc”, “geographic location (region and locality)”, “region”, “birth_location”, and “INSDC center name”^10^. In some cases, (n = 5,710, 27% of sequences), no geographical data was assigned due to lack of metadata. World map and PV location was visualized in R, using packages *ggplot2* (v4.0.0) *raster* (v3.6-32), *rnaturalearth* and *rnaturalearthdata* (v1.1.0) and *sf* (v1.0-21) [https://github.com/syueqiao/pvs_in_logan]^84–87^.

### Case Study Papillomaviruses

To select PVs for case study, sequences from different regions of the phylogenetic tree, long contig length, novelty, and personal interest were prioritized. These examples are not meant to comprehensively characterize all PVs in the tree, but rather a subset that captures the breadth of discovery. To analyze structures of L1 proteins for the PV case studies, each stop-stop ORF was folded in the Alphafold 3 webserver [.cif files available at https://github.com/syueqiao/pvs_in_logan/files/structures]^88^. Structures were rendered in PyMOL (v3.1.6.1), using custom colors to visualize plDDT scores^38,89^. Alignment to HPV 16 L1 (PDB:1DZL) was performed in PyMOL with default parameters^37^. Predicted aligned error plots were generated in R with *ggplot2* (v4.0.0) and *jsonlite* (v2.0.0)^84,90^. EMBOSS *getorf*, DIAMOND *blastp* [*nr* access date 2026-02-05] and InterProScan webserver were used to retrieve and confirm the identity of annotated ORFs^32,36,69^. ORFs for gene graphs were visualized with *gggenes* (v0.5.1) and *ggplot2* (v4.0.0)^84,91^. Subtrees were manually selected and visualized using *ggtree* (v3.12.0) and *treeio* (v1.28.0), between 1 – 5 levels back from the tip of interest, depending on the complexity of the surrounding branches^80,92^. Map subsets were generated using *raster* (v3.6-32), *geodata* (v0.6-6), and *tidyterra* (v0.7.2)^85,93,94^. The most precise location for each sequence available was used, based on manual interrogation of metadata or publications. Two representative contigs, associated with white-bellied pangolin and legless lizard, were chosen as exemplars within the main text.

### Papillomavirus Biomes

All geographical analysis and visualizations were carried out in R, using packages *cowplot* (v1.2.0), *dbscan* (v1.2.3), *ggplot2* (v4.0.0), *ggpmisc* (v0.6.2), *ggrepel* (v0.9.6), *jsonlite* (v2.0.0), *rnaturalearth* (v1.1.0), *rnaturalearthdata* (v1.0.0), *sf* (v1.0-21), *terra* (v1.8-80), *countries* (v.1.2.2), *RColorBrewer* (v1.1-3), *extrafont* (v0.20), and *ggnewscale* (v0.5.2)^84,86,87,90,93,95–103^.

To investigate the distribution of PVs and their sampling throughout the SRA, an equal-area, pseudocylindrical project of the world map (Mollweide, ESRI:54009) was overlaid by a 50km-by-50km grid (n = 205,409 squares)^104^. Each grid was assigned a biome category, based on (in order of priority) the 2017 World Wildlife Fund (WWF) Ecoregions and Natural Earth datasets (Oceans, Rivers and Lake Centerlines, and Lakes and Reservoirs, at 1:50m)^86,87,105^. This was accomplished by generating centroids for each of the grids with *st_centroid* and finding the intersection of this centroid with a biome category. If no biome was assigned, the grid square was considered “Terrestrial Other”, which corresponds to areas of rock and ice. Each biome was assigned a distinct color for visualization. The number of SRA runs attributed to locations within a grid square was counted to approximate ‘sampling effort’ in that square.

Each SRA-derived PV was assigned an associated biome based on the location derived from the BioSample identifier. For visualization of PVs and biomes, a 50-100km buffer of the grid was used to increase visual clarity. To identify geographic squares that are enriched for novel PV types, each grid square was randomly assigned a novel-to-known-type ratio from the dataset without replacement, for 2000 permutations, to generate a standard deviation and mean ratio for every square. A Z-score for each square was generated ([Actual ratio - permuted mean ratio]/standard deviation of permuted ratio) and the corresponding *p*-value calculated with *pnorm* in R. Squares with *p* < 0.05 were considered significantly enriched for novel PV types. No multiple testing correction was applied.

To determine if grids of high novel type proportion were spatially clustered, the nearest eight neighbour grids were used to calculate spatial weights. Moran’s I was used to examine the global distribution, based on the previously generated Z-scores, and Getis-Ord Gi* to identify grids with values driving grouping^33–35^. To visualize these relationships, grids with novel PV types were clustered with a 1000km gap tolerance, and a 500km buffer was used to represent broad regions of high novel type proportion as ‘hotspots’. No significant ‘cold spots’ of PV novelty were identified.

## DATA AVAILABILITY

The analysis code, sequences, and data for this study are freely available. Analysis code and scripts are in https://github.com/syueqiao/pvs_in_logan. Supplementary Data files and sequences are in https://zenodo.org/records/19672425.

## ACKNOWLEDGEMENTS

We would like to thank R. C. Edgar, R. Connor, A. Morales-Tapia, H. Debat, F. Mostefai, K. Rothe, and E. Park for their helpful discussion and comments on the manuscript.

We are grateful to the entire team managing the NCBI SRA/EBI ENA and the biology community for openly sharing their data. Computing resources were provided by the University of Toronto Cloud Research Lab at The Donnelly, powered by AWS. A.B was supported by Canadian Institutes for Health Research (CIHR) project grant PTJ-496709, and as a Canadian Institute for Advanced Research (CIFAR) Global Azrieli Scholar of the CIFAR Fungal Kingdom: Threats & Opportunities program. R.C was supported by Agence Nationale de la Recherche (ANR) grants ANR-22-CE45-0007, ANR-19-CE45-0008, PIA/ANR16-CONV-0005, ANR-19-P3IA-0001, ANR-21-CE46-0012-03, and Horizon Europe grants No. 872539, 956229, 101047160 and 101088572. J.S was supported by the Natural Sciences and Engineering Research Council of Canada (NSERC CGS-M).

## SUPPLEMENTARY FIGURES

**Supplementary Figure S1.**
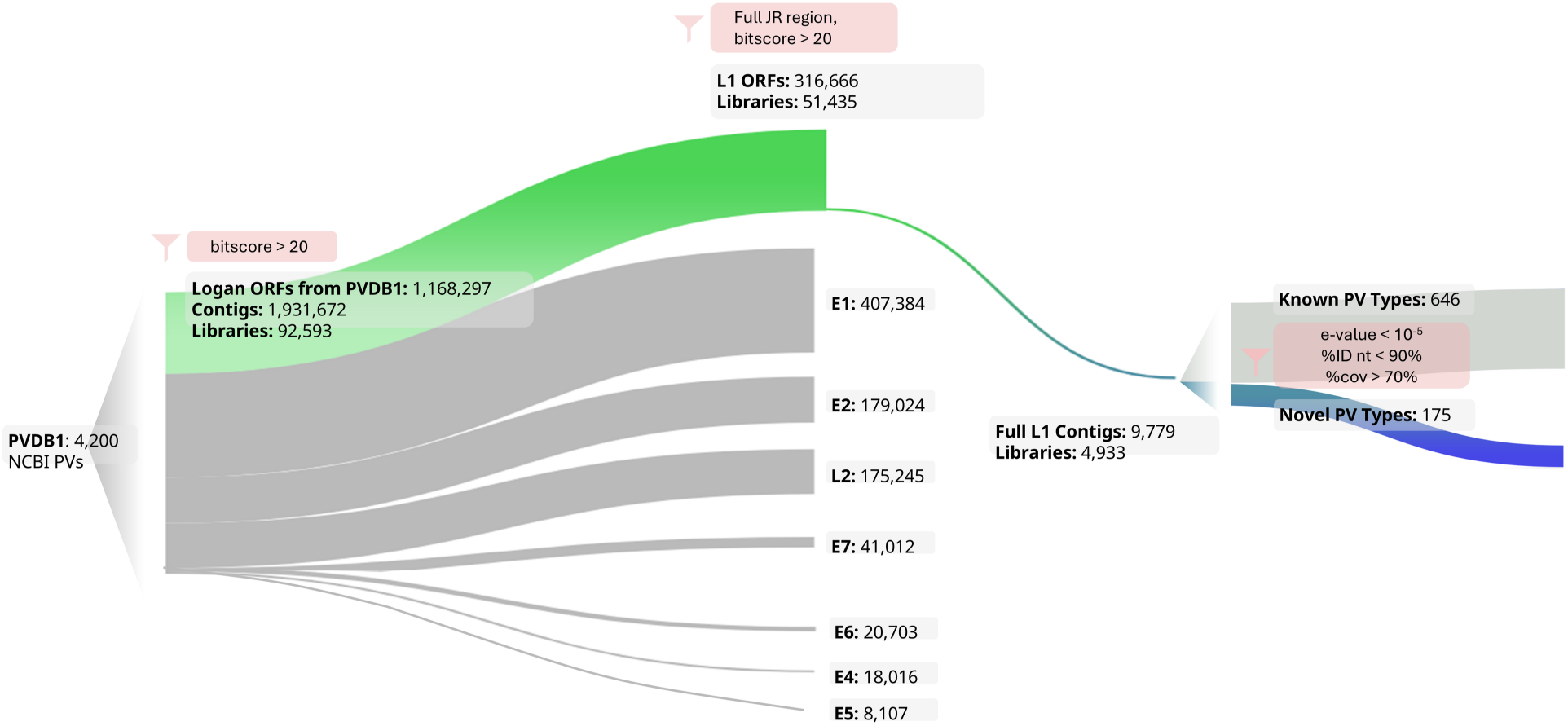
Flowchart visualizing first round of PV searching, using PVDB1, and *Logan* v1.0.

**Supplementary Figure S2.**
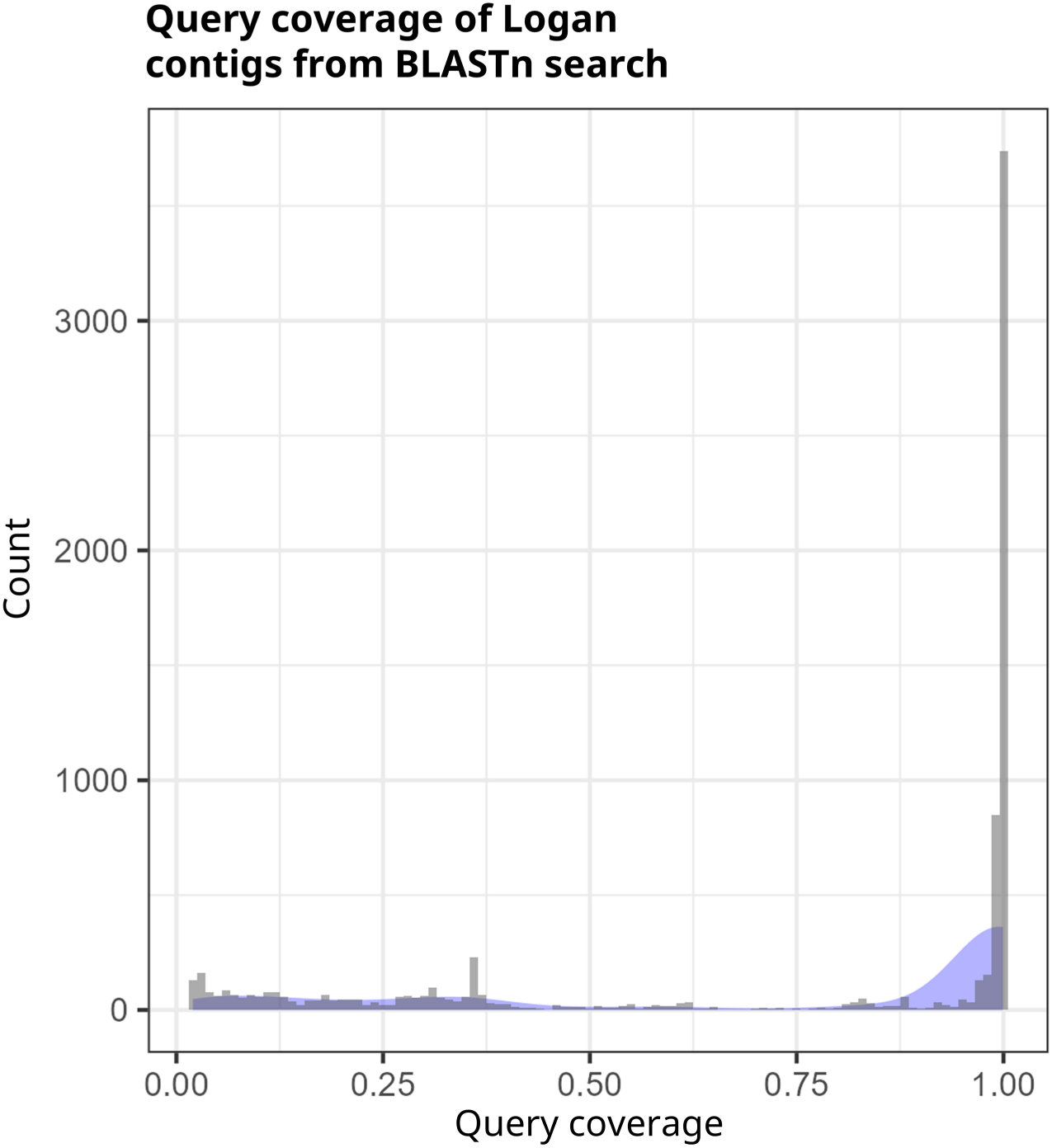
Distribution of query coverage following *blastn* search of *Logan* L1 contigs. Query coverage was calculated by dividing the matched residues over the total length of the input contig.

**Supplementary Figure S3.**
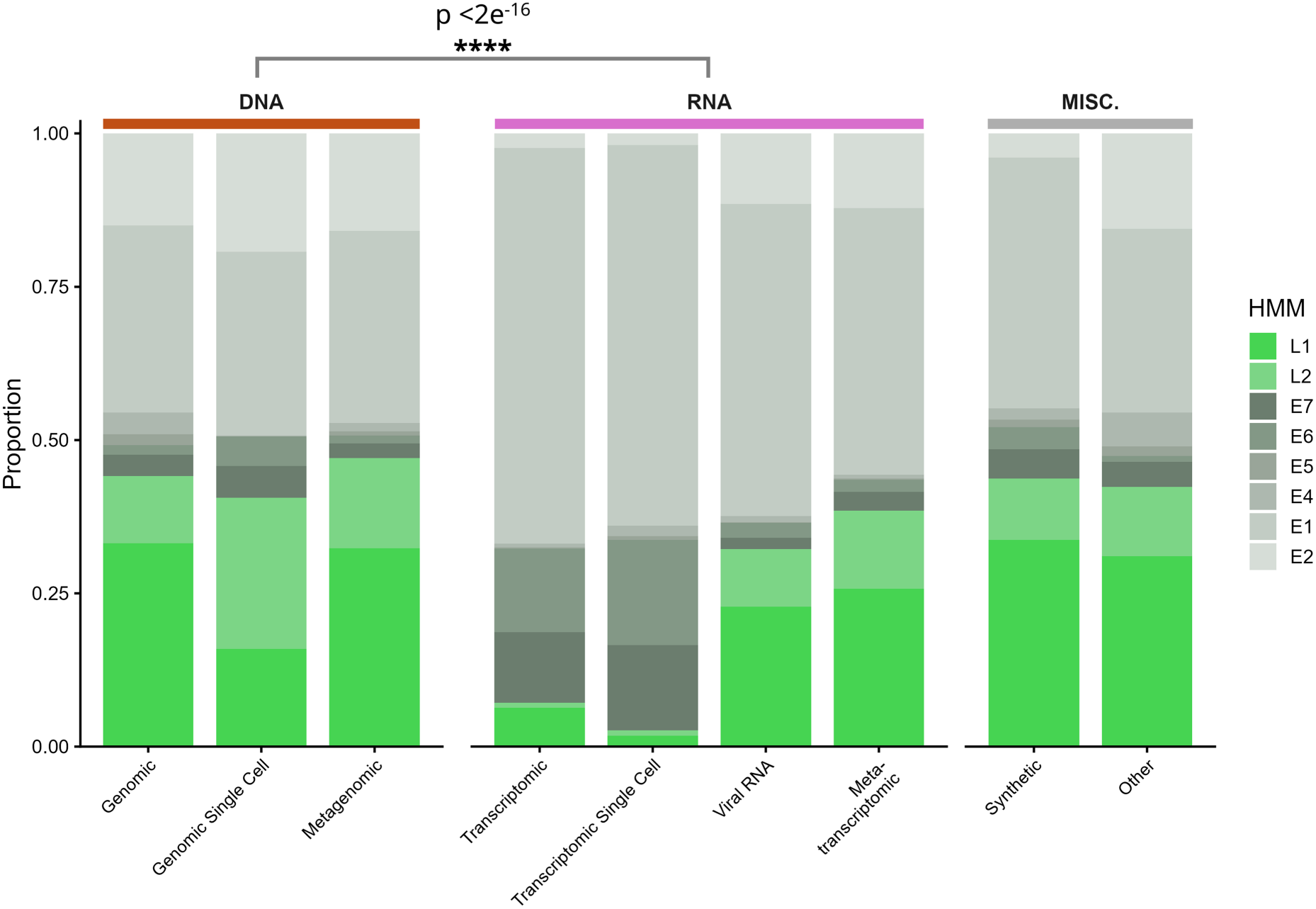
Proportion of PV gene coverage by SRA library type. Stacked bars show the relative proportion of PV gene annotations (L1, L2, E1–E7), weighted by *ka.f* (k-mer-based coverage), across SRA library source categories grouped into DNA-, RNA-based, and miscellaneous. For multi-domain genes, *ka.f* values were averaged across E1 (E1_N, E1_DBD, E1_C) and E2 (E2_N, E2_C) sub-profiles. Chi-squared test was carried out between E and L ratios in pooled DNA and RNA sources.

**Supplementary Figure S4.**
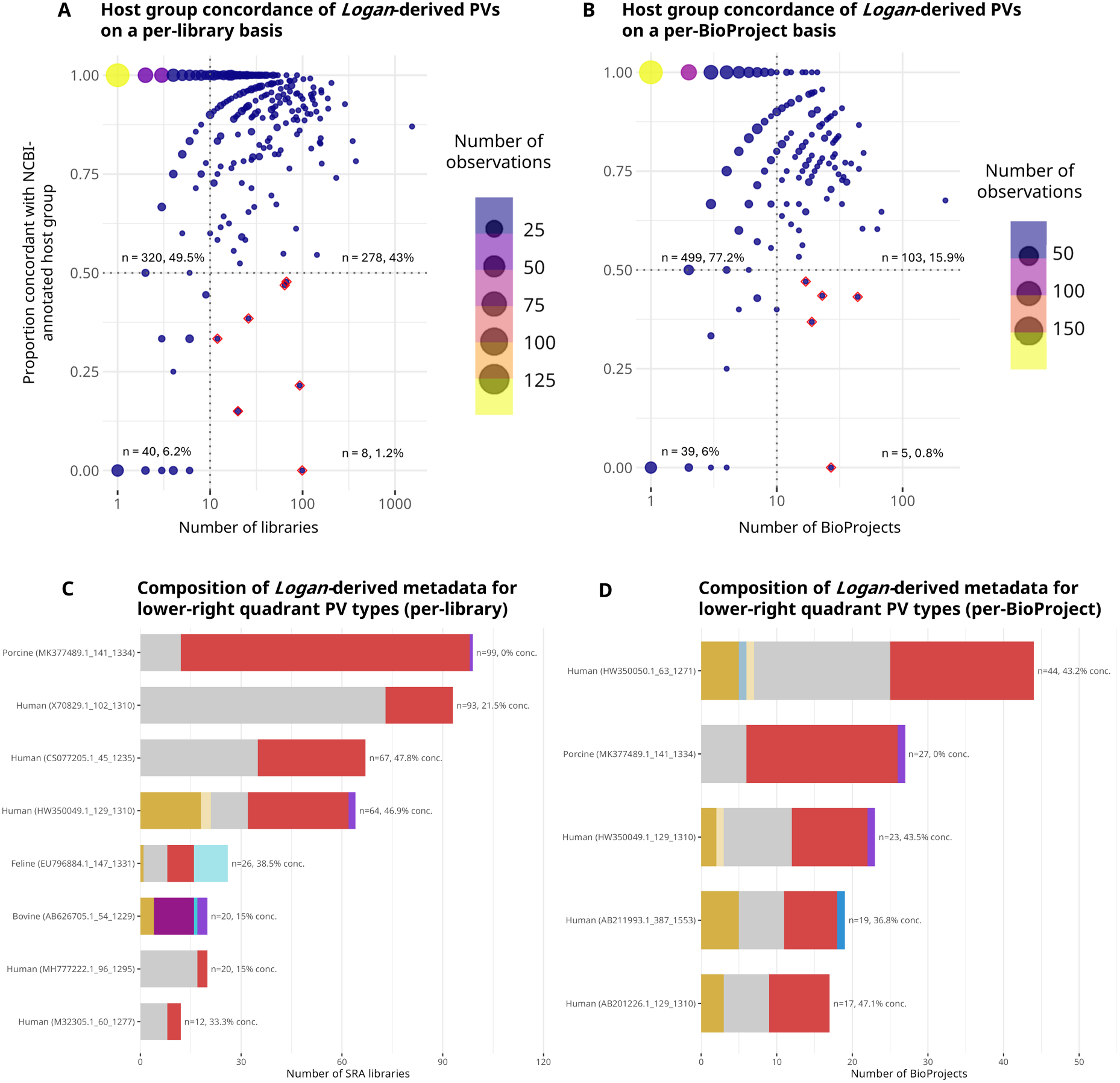
Host group concordance of metadata-annotated PVs from *Logan* in comparison to NCBI annotations. Each point represents one of the 646 NCBI Virus PV types found in *Logan*. The size and color represent the frequency at which the specific percent concordance and number of **A.** libraries or **B.** BioProjects were observed. The exact number and overall percent of types falling into each quadrant is indicated. Points with red diamond outlines indicate types in which more than 10 observations were made, but concordance was lower than expected. **C, D. Distribution of associations derived from *Logan* metadata, where matches to NCBI Virus were found.** On both a per-library and per-BioProject basis, the metadata-derived host for sequences with many observations but low concordance was visualized.

**Supplementary Figure S5.**
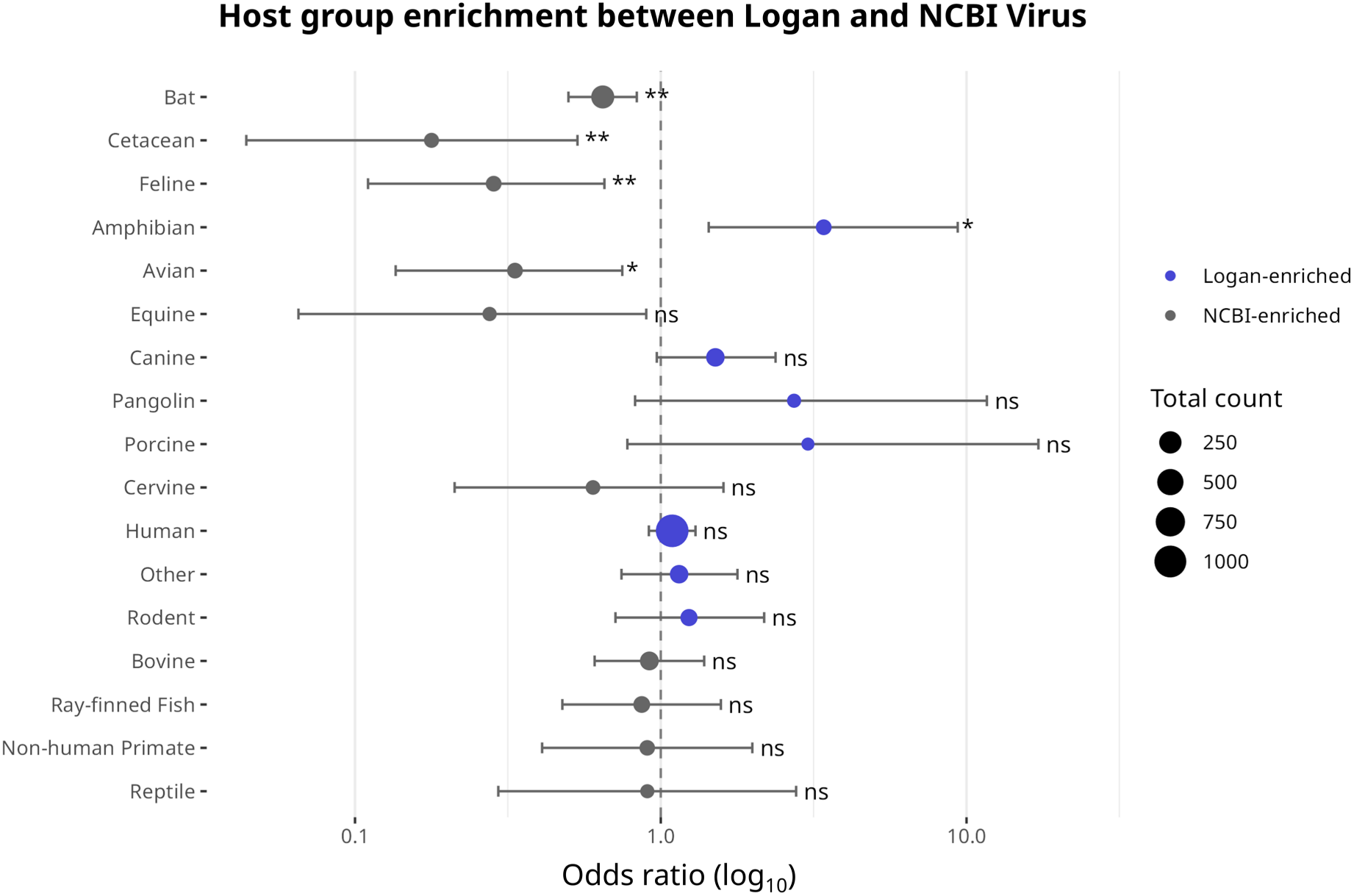
Host group enrichment analysis comparing *Logan* and NCBI Virus PVs. Odds ratios (log scale) from Fisher’s exact tests comparing the proportion of each host group between PV types in *Logan* (n = 1,097) and NCBI Virus (n = 992). Odds ratios >1 (blue) indicate enrichment in Logan; odds ratios <1 (grey) indicate enrichment among NCBI references. Error bars represent 95% confidence intervals. Point size is proportional to the total number of PV types associated with each host group across both datasets. Significance levels are BH-adjusted for multiple comparisons (*p < 0.05, **p < 0.01, ns = not significant).

**Supplementary Figure S6.**
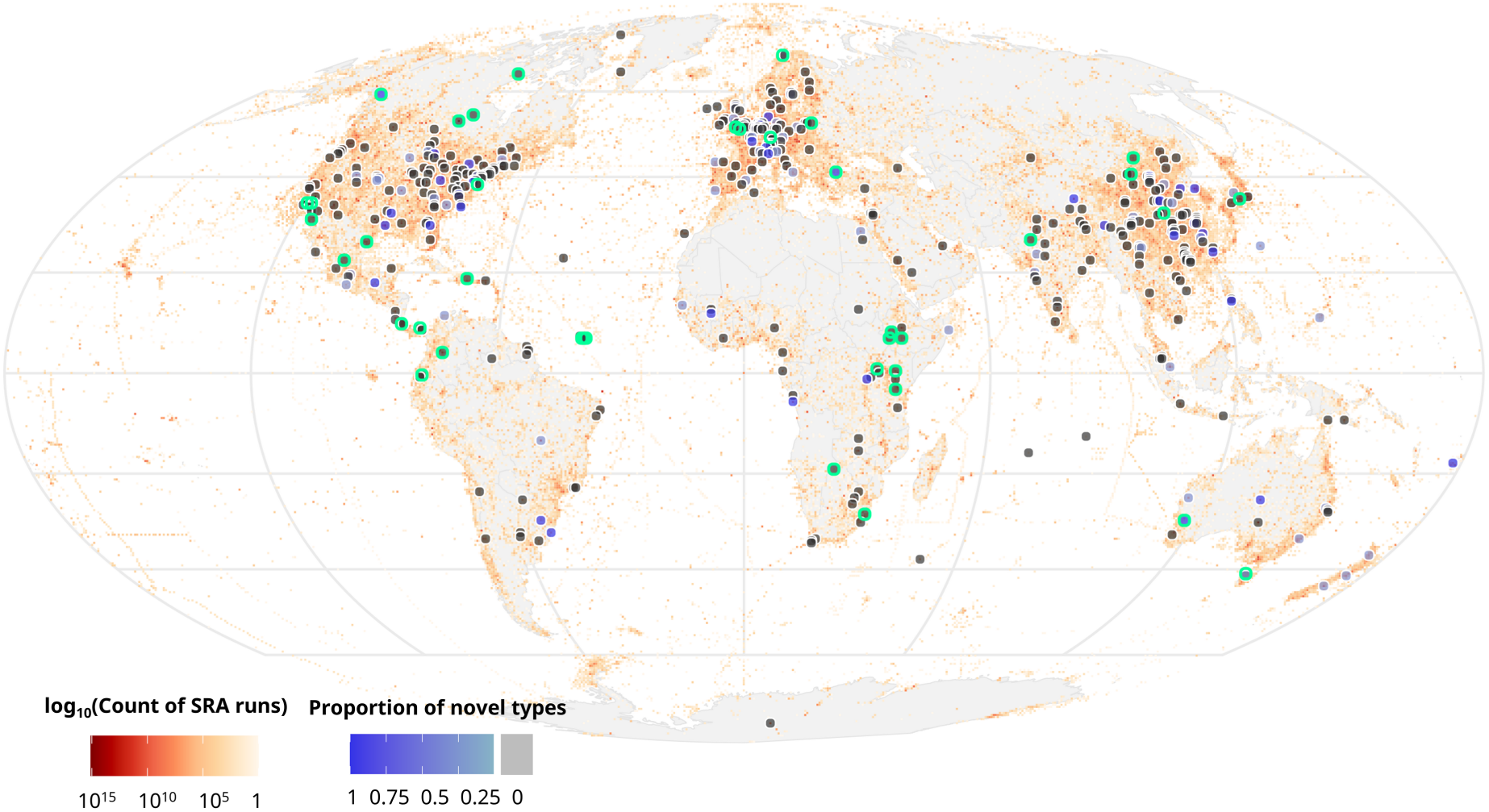
Green outlines of each PV-positive grid square represent those which have statistically significantly more novel PV enrichment following the 2000-iteration permutation test.

**Supplementary Figure S7.**
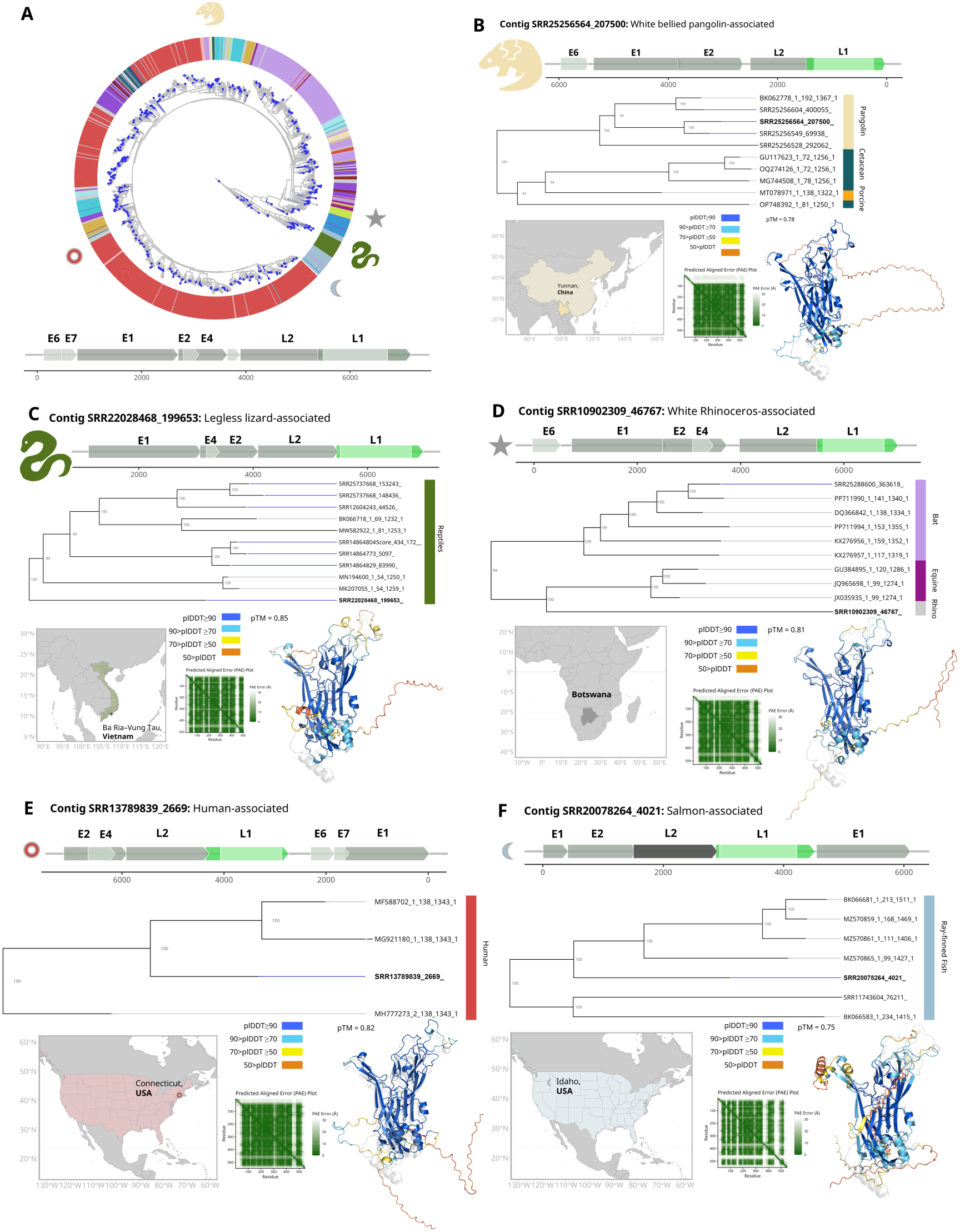
Visualization of all case studies. A) Circular representation of the L1-based papillomavirus phylogenetic tree and generic papillomavirus genome. Blue tips of each branch indicate novel PVs uncovered in this study, and the colored outer ring corresponds to the associated host group. Generic PV genome was visualized using the expected average size of each ORF to show the consistent synteny throughout all case studies.

**Supplementary Figure S8.**
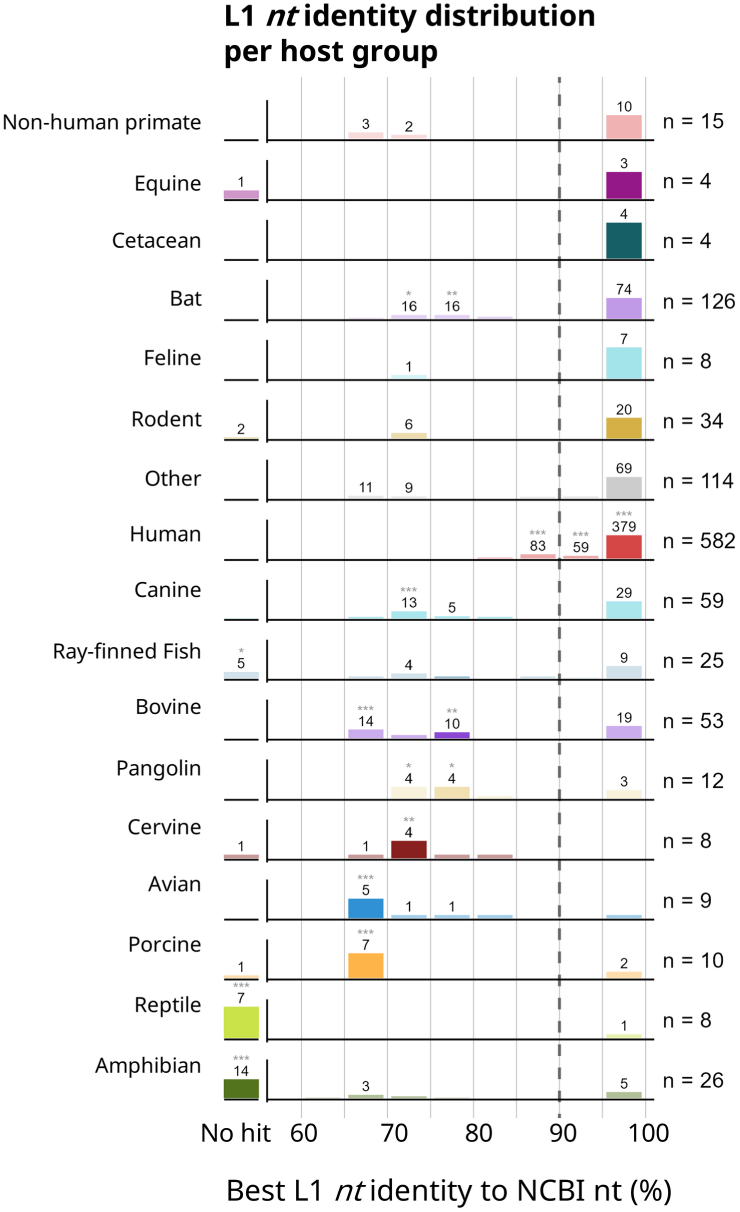
Best-hit *Logan* L1 nucleotide identities per host group. Bars show the number of centroid clusters (n = 1,097), binned by 5% increments, and the percent nucleotide identity to the best hit in the *nt* database with at least 70% query coverage. Centroids with no hit are shown in the ‘no hit’ bin, and the dashed line marks the 90% species-level demarcation. The median bin for each host is colored in full saturation, and asterisks mark bins which are over-represented relative to overall frequency (Fisher’s exact test, Benjamini–Hochberg adjusted; * p < 0.05, ** p < 0.01, *** p < 0.001). Totals per host are annotated to the right.

**Supplementary Table S1.** Distribution of known and novel PV types across host organism groups. Count of unique PV types associated with each host organism group, based on the sequences used for the phylogenetic analysis. Host groups were manually assigned from SRA metadata or PaVE annotation. Groups with fewer than nine total types are combined under the ‘Other’ category.

| host_group | logan_pos | logan_neg | ncbi_pos | ncbi_neg | p_value | odds_ratio | p_adj |
| --- | --- | --- | --- | --- | --- | --- | --- |
| Bat | 126 | 972 | 166 | 826 | 0.000629 | 1.549978 | 0.004405 |
| Cetacean | 4 | 1094 | 20 | 972 | 0.000349 | 5.623248 | 0.004405 |
| Feline | 8 | 1090 | 25 | 967 | 0.00124 | 3.520539 | 0.005788 |
| Amphibian | 26 | 1072 | 7 | 985 | 0.002411 | 0.293182 | 0.00844 |
| Avian | 9 | 1089 | 24 | 968 | 0.004305 | 2.998517 | 0.012054 |
| Equine | 4 | 1094 | 13 | 979 | 0.025283 | 3.629853 | 0.058994 |
| Canine | 59 | 1039 | 36 | 956 | 0.058795 | 0.663275 | 0.117591 |
| Cervine | 8 | 1090 | 12 | 980 | 0.271127 | 1.667945 | 0.474472 |
| Human | 583 | 515 | 505 | 487 | 0.334807 | 0.916051 | 0.52081 |
| Other | 53 | 1045 | 42 | 950 | 0.529985 | 0.871756 | 0.674527 |
| Rodent | 34 | 1064 | 25 | 967 | 0.508826 | 0.809134 | 0.674527 |
| Bovine | 53 | 1045 | 52 | 940 | 0.689137 | 1.09068 | 0.742147 |
| Ray-finned Fish | 25 | 1073 | 26 | 966 | 0.671036 | 1.155115 | 0.742147 |
| Non-human Primate | 15 | 1083 | 15 | 977 | 0.854737 | 1.10844 | 0.854737 |

## SUPPLEMENTARY FILES

*Available at* https://zenodo.org/records/19672425

**Supplementary File 1. PaVE L1 alignment.** Aligned L1 genes retrieved from the PaVE webportal, used to generate a multiple sequence alignment^27^.

**Supplementary File 2. NCBI_Virus_PV_ORFs_trimmed.** BED-file for trimming ORFs from NCBI Virus used for the initial search of PVs in Logan, used to generate PVDB1 and PVDB2.

**Supplementary File 3. jrHMM.** Custom jrHMM, built using the L1 sequences from NCBI Virus, PaVE, and PVDB1, for each jelly roll region. Sequences were clustered at 90% amino acid identity prior to alignment.

**Supplementary File 4. Logan_Outputs_R1.** FASTA-file containing output contigs of the first Logan search with PVDB1.

**Supplementary File 5. Logan_Outputs_R2.** FASTA-file containing output contigs of the second Logan search with PVDB2.

**Supplementary File 6. Host_Assignments_Generalization.** Putative assignments of host groups for each representative sequence, at both a broad and species level, when information was available.

**Supplementary File 7. All_PVs_Metadata.** Comprehensive table for all PVs from this study and NCBI Virus, including accessions, original metadata, generalized host groups, geographic, and biome information where available.

**Supplementary File 8. Host_Concordance_Full_Table.** Table containing data used for host concordance analysis.

**Supplementary File 9. BLASTn_search_identity.** Full tab-separated table of *blastn* results.

**Supplementary File 10. PV_Tree_Seqs.** Multiple sequence alignment file used to generate a comprehensive phylogenetic tree. Contains both novel and known PVs from Logan and NCBI Virus respectively.

**Supplementary File 11. NCBI_Only_Seqs.** Table containing accessions of NCBI Virus PVs not found in *Logan*.

**Supplementary File 12. BLASTp_hits.** Full tab-separated table of *blastp* results of case study gene maps.

